# Widespread Genomic Islands in Giant Viruses Shape Genome Plasticity and Mosaicism

**DOI:** 10.64898/2026.04.13.718226

**Authors:** Benjamin Minch, Mohammad Moniruzzaman

## Abstract

Giant viruses in the phylum *Nucleocytoviricota* possess exceptionally large and mosaic genomes, yet the mechanisms underlying their remarkable genomic plasticity remain poorly understood. Genomic islands are large dynamic genomic regions that are major drivers of genome diversification and adaptation in bacteria. However, their contribution to genome evolution in giant viruses remains largely unexplored. Here, we systematically characterize the genomic island landscape of giant viruses using 369 high-quality genomes spanning cultured isolates and long-read metagenome-assembled genomes. We identify 307 genomic islands across >50% of the genomes, demonstrating that these regions are pervasive across *Nucleocytoviricota* diversity. Giant virus genomic islands are frequently associated with genomic hypervariability and enriched in genes involved in host interaction, particularly surface adhesion proteins, suggesting key roles in host adaptation during virus-host arms race. Comparative analyses further reveal these islands to be major hotspots of genome diversification, exhibiting frequent gain, loss, and rearrangement even among highly similar genomes, with evidence that entire island regions can be exchanged among closely related viral populations. Notably, 37% of genomic islands are enriched in bacterial homologs, and several exhibit striking synteny with genomic regions recovered from environmentally co-occurring bacterial genomes, supporting large-scale genetic exchange between bacteria and giant viruses. Together, these findings identify genomic islands as pervasive and dynamic drivers of giant virus genome evolution, providing a mechanistic framework for genome plasticity, mosaicism, and adaptive potential of giant viruses.

## Introduction

Giant viruses in the phylum *Nucleocytoviricota* (NCVs) are a unique group of eukaryote-infecting viruses with both large capsid (up to 2um) and genome sizes (up to 2.5 Mbp) [1], [2], [3], [4]. These viruses have redefined what was previously the limits of virus functional potential, as they have the ability to encode many genes previously unknown in the virosphere [3], [5]. There are currently six identified orders of NCVs – *Imitervirales, Algavirales, Pimascovirales, Asfuvirales, Pandoravirales*, and *Chitovirales* [6]. These viruses are widespread in the global oceans, lakes, and sediments and have a seemingly broad host range, consisting mainly of single-celled eukaryotes or protists [4], [7]. NCVs infecting protists have the potential to have significant impact on planetary biogeochemical cycles due to both the large biomass of protists on the planet and the known potential of NCVs to modulate host metabolism and carbon fates [5], [8], [9].

While their ecological significance has been well established, many questions remain regarding the large size of NCV genomes, as they are stand-outs in the virosphere [10], [11]. Most viruses exist as largely successful replicators due to their small genome size, allowing for quick and efficient replication [12], [13]. Even viruses infecting complex organisms such as mammals have quite streamlined genomic resources available to them [14], [15]. This is not the case for NCVs, as their genomes are often larger than 200 kilobase pairs (kbp) and contain many proteins that do not seem immediately useful for viral replication [3], [16]. This gigantism of NCV genomes has evolved multiple times throughout their evolutionary history, and it is thought that the pattern of gene gain and loss follows an “accordion model”[17], [18]. These gene gains are thought to be primarily an equal contribution of gene transfer from other viruses (vHGT), gene duplications, and originations [19].

On top of having large genomes, NCV genomes are often seen as mosaic, comprising many genes from diverse cellular lineages as well as a large number of ORFan genes [2], [20], [21]. The large amount of genetic material from cellular sources within NCV genomes initially led researchers to hypothesize that these viruses derived from an ancient cellular ancestor, an idea that has since lost favor [11], [22], [23]. While we know that NCVs acquire many genes through horizontal gene transfer (HGT), we still lack a clear understanding of the genomic constraints and ecological interactions facilitating these acquisitions.

From extensive research on HGT in bacteria, we know that gene exchange can occur on the individual level as well as in large chunks in the form of genomic islands [24-26]. Genomic islands (GIs) are well characterized in bacteria as large segments of DNA often harboring many horizontally transferred genes [24], [25]. These regions are also usually distinct in nucleotide composition signatures compared to the surrounding genome and are enriched in genes involved in pathogenesis, symbiosis, antibiotic resistance, or response to different environmental factors [24], [26]. GIs often play a crucial role in the evolution and adaptability of bacteria as they introduce genetic diversity and improve fitness in changing environments [25]. GIs are typically fast evolving, with higher mutation rates and hypervariability than the surrounding genome. In this context, hypervariability refers to the fact that rapid gain or loss in genes and fast evolving gene compendium makes islands within closely-related bacterial strains slightly different from each other[24], [27].

Since genomic islands (10-200 kbp) [28] are often as large or larger than many viral genomes, we often don’t associate them with viral populations. The discovery of NCVs, however, calls for a reevaluation of the role of genomic islands in viruses, as these viruses have sufficiently large genomes to contain island regions. Since their discovery in 2003, there have been a few examples of putative hypervariable regions found within a few cultured NCV genomes [29], [30], [31], [32], as well as metagenomic islands (regions of the genome with anomalous metagenomic read recruitment) in uncultured genomes [33], suggesting that regions with features characteristic of genomic islands exist in NCVs. However,their distribution across NCV phylogeny, functional characteristics and ecological significance in NCVs have largely remained elusive. Part of this is due to the majority of NCV genomes coming from short-read, metagenomic binning of NCVs [3], [4], [34], which is a barrier against assembling NCV genomes with high contiguity. However, recent developments in long-read sequencing represents a significant advance in reducing this barrier given the possibility that this technology can uncover complete or near-complete genomes from metagenomic datasets.

Given that there is little information on the origin and functional composition of these islands in NCVs, a method is necessary to identify them that is function agnostic but still rooted in the fundamental nature of GIs - such as deviated nucleotide composition. Motivated by this, we developed a tool that can identify these regions in NCVs based on compositional deviation from the surrounding genomic context. Here, we apply this tool on a curated set of long-read and cultured representative NCV genomes to uncover the genomic island landscape in NCVs [35], [36]. We demonstrate that genomic islands are widespread among all orders of *Nucleocytoviricota* and are enriched in surface adhesion, replication, and metabolic genes, providing evidence of their use as host-adaptation mechanisms. Additionally, leveraging long-read genomic data, we show hypervariability of these regions within closely related strains, possible HGT of these islands across strains, and their contribution to genomic plasticity of closely related NCVs. Furthermore, we show that many genomic islands are enriched in genes with bacterial homologs, indicating a strong bacterial contribution to their gene content. Notably, in several cases these islands exhibit remarkable synteny and gene content similarity to genomic regions of bacteria that co-occur within the same environmental sample, providing evidence of large-scale transfer of genomic regions between bacteria and NCVs shaping the GI landscape in NCVs. Taken together, these findings position genomic islands as key mediators through which NCVs acquire adaptive traits, shaping their ecological interactions, genome evolution and host-virus arms race. The results also lay the groundwork for future research into the ecological roles of genomic islands in NCVs and how they may be shaping the ecology and evolution of these viruses on a global scale.

## Materials and Methods

### Gathering high-quality NCV genomes

To identify genomic islands across the diversity of NCVs, we obtained NCV genomes that were either complete or near-complete (based on the presence of key hallmark genes) from various sources. These included 59 genomes from cultured viruses [6] and 146 published nanopore long-read genomes from Lake Biwa, Japan [35]. In addition to these, we identified an additional 164 single-contig nanopore long-read genomes from published data from the south bay of the San Francisco Estuary [36] using BEREN in contig mode [37].

All of these genomes were selected based on the presence of at least 5 Nucleocytoviricota marker genes [6], identified using the ncldv_markersearch tool [6] (Table S1). Among the 5 marker genes, genomes had to contain a DEAD/SNF2-like helicase (SNFII), DNA polymerase B (polB), Packaging ATPase (A32), and Poxvirus Late Transcription Factor VLTF3 (VLTF3), as these are common genes used in establishing the phylogeny and identity of NCVs [4], [6]. This resulted in a total of 369 high-quality or near-complete genomes for downstream analysis. A concatenated phylogeny of these 4 marker genes was created using MAFFT [38] for alignment, trimAl with ‘-gt 0.1’ [39] for alignment trimming, and IQTREE (LG+F+R10 model) [40]. The resulting tree was visualized using Anvi’o [41] (Figure 1).

**Figure 1.**
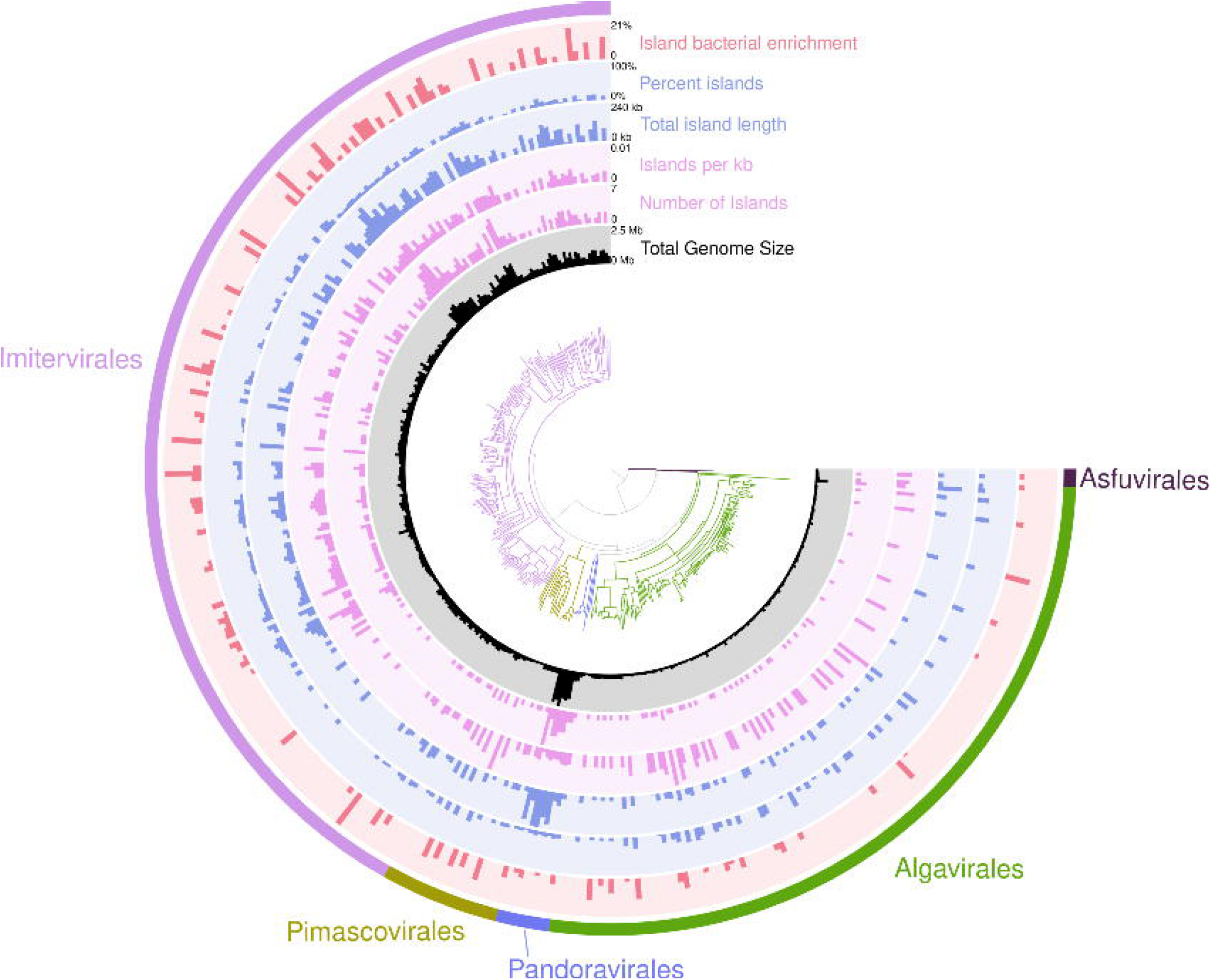
Distribution of genomic islands in giant viruses. A phylogenetic tree using a concatenated alignment of DEAD/SNF2-like helicase (SNFII), DNA polymerase B (polB), Packaging ATPase (A32), and Poxvirus Late Transcription Factor VLTF3 (VLTF3) was constructed using our 369 reference genomes across 5 orders of *Nucleocytoviricota*. Rings around the tree are as follows: (1) Total genome size, (2) the total number of islands identified, (3) Islands per kbp, normalized to genome size, (4) total length of island regions, (5) percent of the genome that island regions took up, and (6) island bacterial enrichment which is calculated as the difference between the proportion of genes having best hits to bacteria inside and outside the genomic islands in a genome. Negative values were rounded to 0.

### Identification of genomic islands

To identify the genomic islands in the genomes of NCVs, we developed a Python tool (IslaMine) that uses deviations in tetranucleotide frequency (TNF) and flanking direct repeats to identify these regions, common features of genomic islands found in diverse bacteria and eukaryotes [24], [42]. Each genome is first divided into non-overlapping chunks of 5 kbp from which the TNF profile is computed and compared to the TNF of the full genome using a Pearson correlation. The resulting correlation values are converted to a cumulative sum and first derivative to highlight compositional shifts. A sliding window variance is then calculated across the derivative signal to identify regions of local instability. Peaks in the variance signal are then detected using adaptive thresholds based on genome size (90^th^ percentile of the variance), with adjacent or overlapping peaks being merged. An example output can be found in Figure S1a.

Each region of high variance that is predicted through this pipeline is then scanned for the presence of flanking direct repeats (minimum length of 10 bp), +/- 500 bp on either side of the region. Islands and the flanking regions (+-5 kb) were scanned for tRNAs using tRNAscanSE 2.0 [43][24]. The IslaMine tool is publicly available on github (https://github.com/BenMinch/IslaMine).

#### Validation of the tool using known NCV genomic islands

In separate studies, several NCV genomes were reported to contain putative genomic islands [31], [32], We attempted to independently recover these genomic islands using our python tool, IslaMine. Our tool successfully recovered all 21 of the genomic islands present in these genomes, with the identified islands often covering > 90% of the reported island regions (Supplemental Table 1). This validates IslaMine’s ability to identify NCV GIs and indicates that genomic islands previously identified in NCVs share common compositional and structural features (altered tetranucleotide frequency and flanking repeats), which possibly represents generalizable hallmarks of genomic islands in NCV genomes.

#### Identification of metagenomic islands

To identify genomic islands that were also metagenomic islands (regions of a genome with deviations in read mapping coverage), Illumina short-read data from the San Francisco Estuary [36] was mapped to all identified genomes from that region using minimap2 (-pa 95)[44]. Resulting alignments were converted to BAM files with samtools [45] and visualized using a custom Python script to look at coverage across the genome (Figure S1b).

### Functional analysis of genomic islands

For each genomic island, open reading frames (ORFs) were predicted using prodigal-gv [46], [47] and annotated using Annomazing (a HMMER-based annotation pipeline [48]) [34] with the Pfam [49] and GVOG [6] databases (e-value cutoff of 1e-5). Resulting annotations were then categorized based on their Pfam annotation (Structure, Environment Response, Metabolism, Replication, Interaction, or Unknown, (Table S2). Proteins with unknown Pfam domains (DUFs) were additionally annotated using Gaia which utilizes protein embeddings [50]. Mobile genetic elements within the genomic islands were identified using REBASE [51], ISfinder [52], and Pfam databases [49].

To determine genes enriched in islands of specific phylogenetic orders, a Fisher’s exact test was performed utilizing a FDR-corrected p-value of < 0.05 for significance. The same Fisher’s exact test was also used to find genes enriched in island regions when compared to the rest of the genome. Maps of genomic islands were created using lovis4u [53] (Figures 2-3).

**Figure 2.**
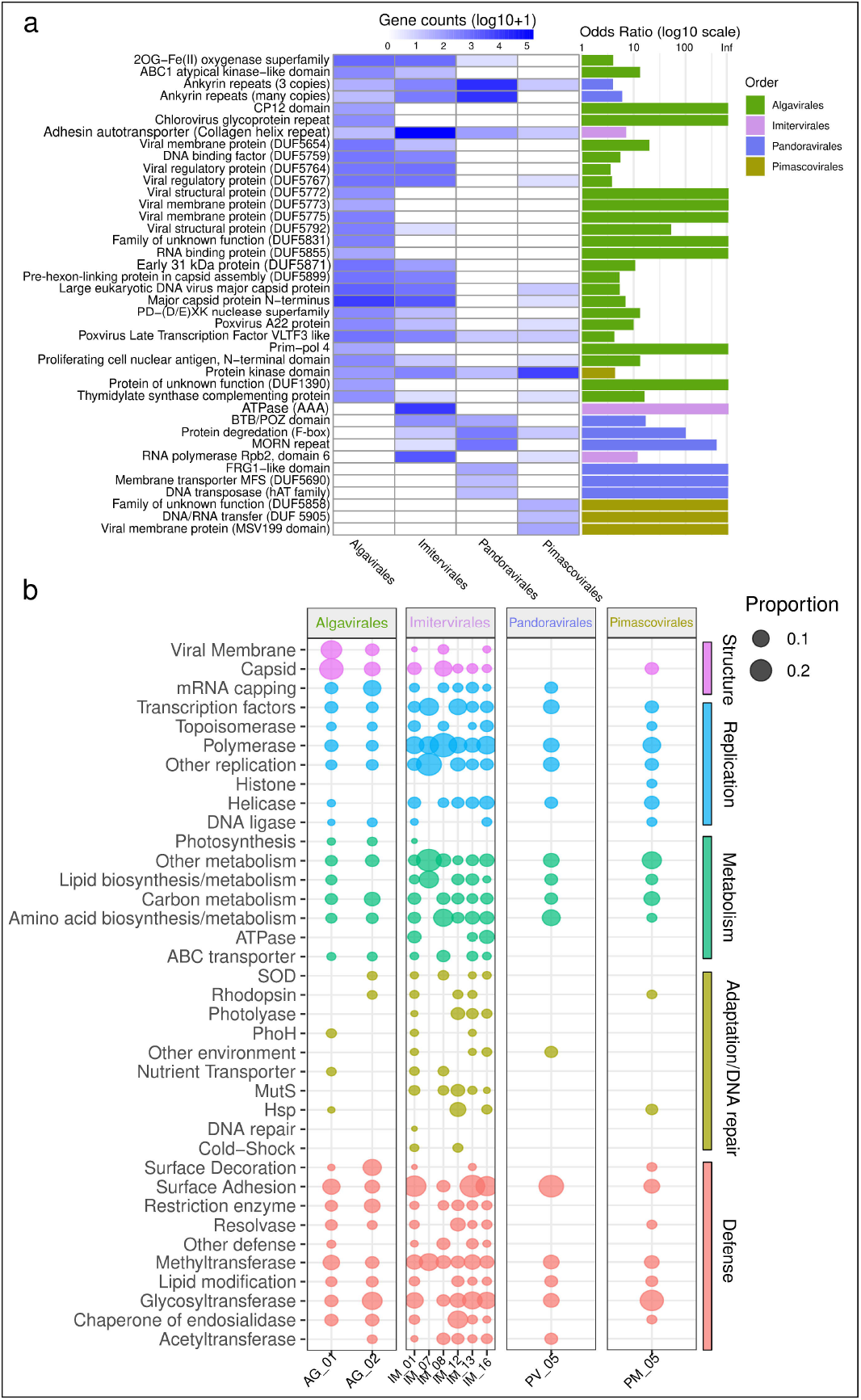
Functional differences in genomic islands across *Nucleocytoviricota* orders. **(a)** A Fisher’s exact test was performed (FDR p-value cutoff of .05), resulting in 42 proteins enriched in islands of a particular order. The heat map shows the log10 gene counts (log10+1 to remove errors), and the barplot shows the odds ratio when comparing that order to all the other orders. **(b)** Proteins of interest associated with viral structure, replication, metabolism, environment response, and interaction were selected based on previous studies [3]. The proportion of islands containing these functions for each *Nucleocytoviricota* family is shown as bubbles in the plot.

**Figure 3.**
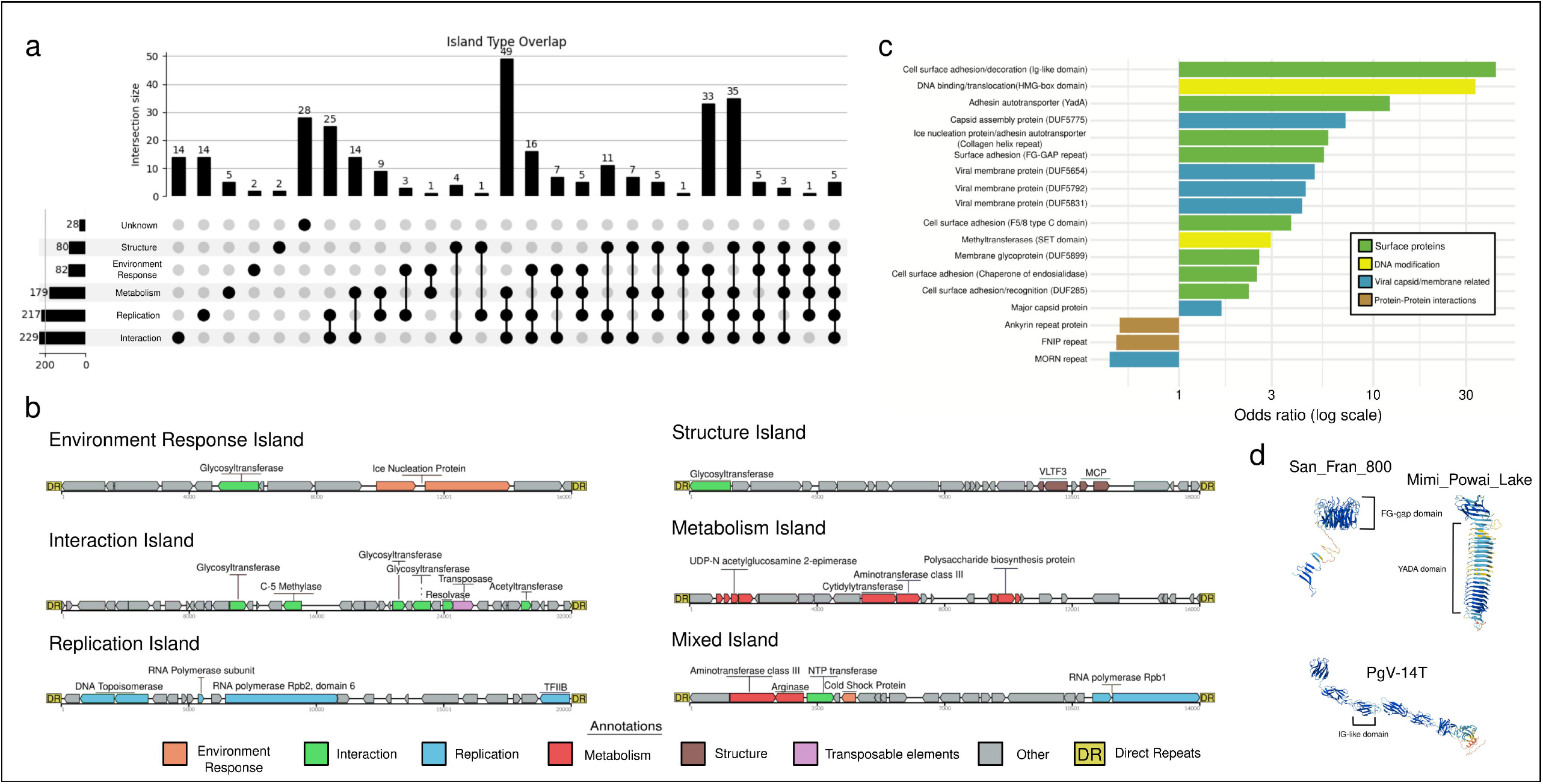
Genomic island functional characteristics. **(a)** An upset plot showing the number of islands encoding different types of functions related to structure, adaptation, metabolism, replication, or host interaction. An island was placed into a category if it encoded at least one gene in that category. Islands entirely composed of genes of unknown function were placed in the unknown category. **(b)** Representative islands for each island type were mapped with lovis4u, highlighting the genes fitting into defined categories. **(c)** A Fisher’s exact test was run comparing all of the proteins inside the genomic islands to the proteins found in giant virus genomes outside the island regions. Enriched proteins (p <0.05) in both groups are shown here with their odds ratios and categorization. **(d)** Representative enriched surface adhesion proteins were folded using alphafold3. The different surface adhesion domains are highlighted.

Proteins encoded within and outside of genomic islands were independently clustered into orthogroups using MMseqs2 (--min-seq-id 0.25 -c 0.5 –cov-mode 1 -s 7.5) [54]. Genomic islands were subsequently clustered functionally based on the presence or absence of orthogroups containing at least five members. To this end, hierarchical clustering was performed using Jaccard distances and the Ward.D2 method implemented in the vegan package [55], [56] in R. The optimal number of clusters was determined by minimizing the Davies-Bouldin index, and the resulting clusters were visualized using pheatmap in R [57].

For certain surface adhesion proteins, phylogenetic trees were built alongside 100 reference sequences from the top 100 hits to the Open Genomes Database (based on embedding distances) [58] within Seqhub [50]. Alignments were made using MAFFT [38] and trimmed using trimAl [39]. Trees were made using IQTREE [40] with 1000 bootstraps and displayed using iTOL [59]. Structures of certain island-encoded proteins were predicted using Alphafold3 [60].

### Locating shared genomic islands from the San Francisco Estuary

To locate similar or shared genomic islands in the co-occurring NCVs in the San Francisco Estuary environmental dataset, we first created a BLAST [61] database of all the genomic islands found from genomes recovered from this location. Additional NCV contigs were recovered from the San Francisco nanopore data utilizing BEREN in ‘contig’ mode [37].

BLASTn was then run with a minimum percent identity of 70% and a minimum coverage of 50% to locate similar islands in these NCV contigs. This analysis revealed a cluster of 56 contigs and genomes that shared a similar genomic island. The BLAST network was visualized using Cytoscape [62] (Figure 4).

**Figure 4.**
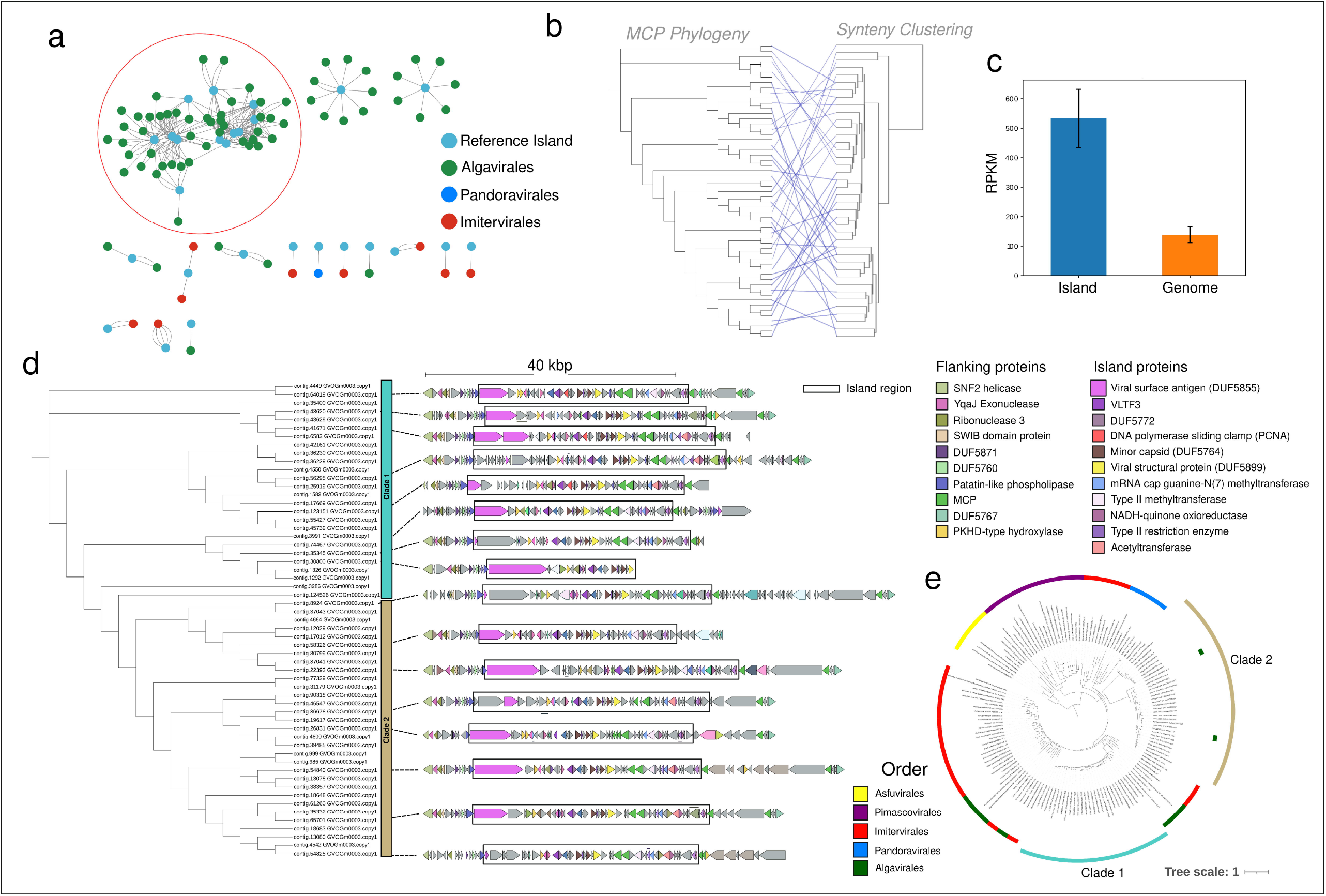
Finding shared genomic islands in the San Francisco Estuary. **(a)** An all-vs-all BLASTn search was performed using reference genomic islands identified in this study and additional giant virus contigs found in the San Francisco Estuary metagenome [36]. The resulting connections at 70% minimum identity and 50% coverage were displayed as a Cytoscape network colored by giant virus order. A large connected node of shared Algavirales islands is highlighted by the red circle. **(b)** A phylogeny of the selected giant viruses with shared island regions was performed using the major capsid protein (MCP) found outside of the island region. Synteny-based hierarchical clustering of the genomic islands was also performed (Figure S8). Lines between islands and MCPs coming from the same genome were drawn to show phylogenetic incongruence. **(c)** Short reads from the San Francisco Estuary dataset were mapped to both the island regions and the other genomic regions of the selected group (red) at 95% identity. The resulting barplot shows the average RPKM of the island and the rest of the genome with error bars representing +/- 2 SEM. **(d)** The same phylogenetic tree as in (b) with overlaid genomic maps of selected contigs showing the annotated conserved island region. **(e)** A broader MCP phylogenetic tree including reference sequences from the GVDB [6]. The two major clades of capsids found within the San Francisco Estuary are highlighted in the outer ring and giant virus phylogenetic order is shown in the inner ring.

To further characterize the distributions of the islands across these contigs, a phylogenetic tree was created for each contig/genome sharing an island using MCP as the phylogenetic marker. MCPs were first aligned with MAFFT [38], then trimmed with trimal ‘-gt 0.1’ [39], and the final tree was constructed using IQTREE [40]. This phylogeny was compared with hierarchical clustering of the gene synteny in the island regions to determine whether gene transfer or the accumulation of mutations was most likely the cause of island heterogeneity [63]. These MCPs were combined with references from the GVDB [6] to create a broader phylogenetic context using the same methods. Additionally, contigs containing shared islands were compared to each other using pyani [64].

To assess island similarity, island proteins were clustered into orthologous groups using proteinortho [65]. The synteny of orthologous groups within islands was compared using Levenshtein distance, treating each protein like a word in a sentence. The resulting synteny dissimilarity was plotted in R.

Finally, to quantify differences in coverage of this widely shared island to the individual NCV contigs and genomes housing the island, short reads from the San Francisco dataset were mapped at 95% identity to both the genomic islands and the rest of the genome individually using minimap2 [44]. The raw read counts were normalized using RPKM.

### Genomic island variability among similar strains

Using a comparative phylogenetic framework, we identified and analyzed prasinoviruses from the San Francisco Estuary to evaluate the prevalence of genomic hypervariability in prasinoviruses. We utilized 19 prasinoviruses from a previous study by Thomy *et al*. [32], as well as 13 new prasinoviruses from the San Francisco Estuary, and created a phylogenetic tree using a concatenated alignment of SFII, PolB, TFIIB, Topoisomerase 2, A32 ATPase, and VLTF3. The alignment was constructed using MAFFT, trimmed with trimAl [38], [39] using ‘-gt 0.1, and the tree was built using IQTREE [40]. Maps for all the genomic islands in the prasinoviruses were created using lovis4u [53] (Figure 5a).

**Figure 5.**
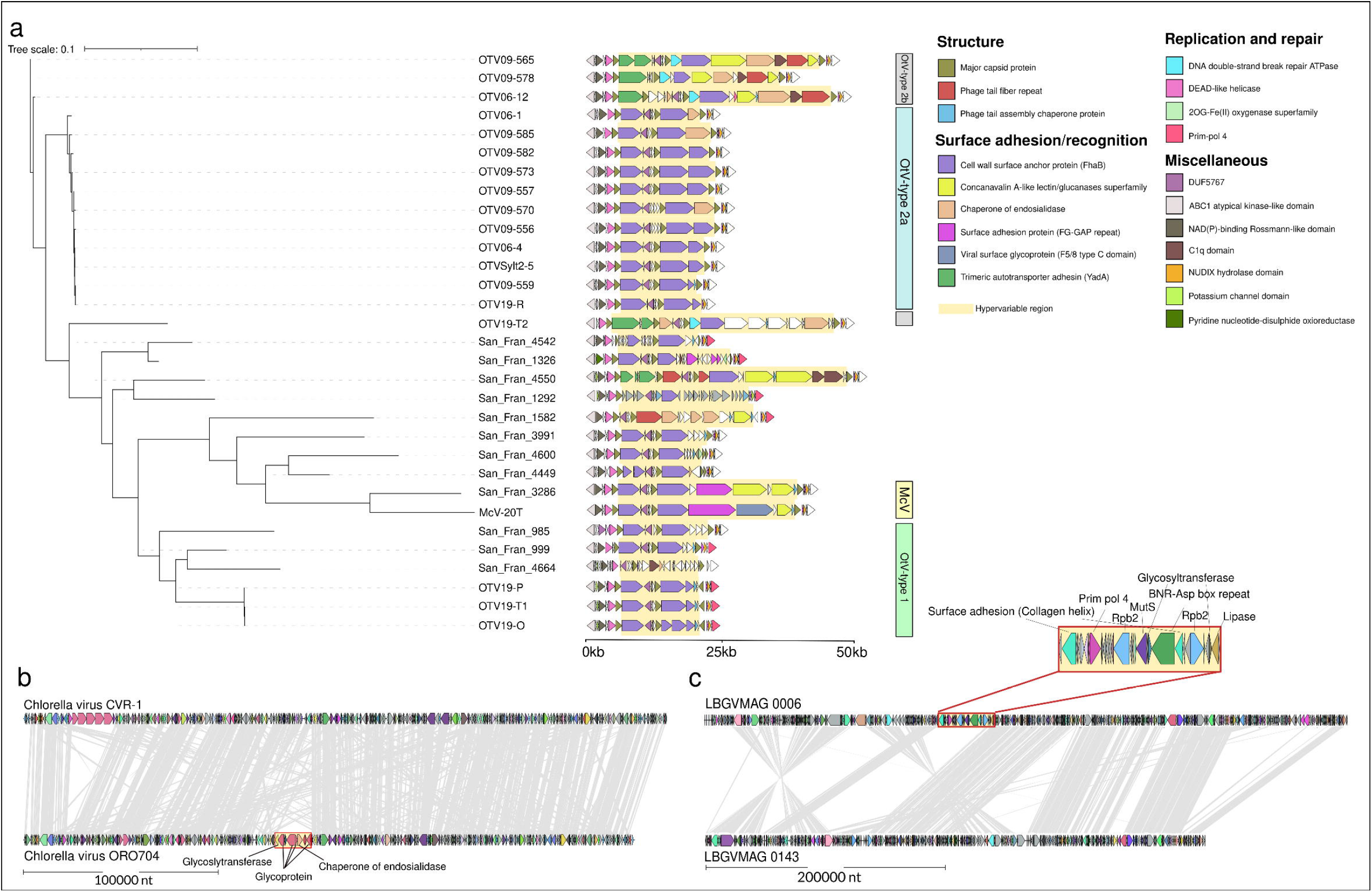
Genomic island variability across similar genomes. **(a)** A phylogenetic tree consisting of a concatenated alignment of SFII, PolB, TFIIB, Topoisomerase 2, A32 ATPase, and VLTF3 was constructed using reference prasinoviruses from Thomy et al. (2026) [32] and newly discovered prasinoviruses from the San Francisco Estuary data in this study. The hypervariable region identified in Thomy et al. (2026) is highlighted in yellow along with gene functional annotations and phylogenetic groups. **(b**,**c)** Complete genome plots of two pairs of similar viruses with different genomic islands are shown with their island regions highlighted in yellow. Vertical grey bars represent synteny of 90% identity or greater at the amino acid level. All plots were made using lovis4u.

Additional pairs of similar Mimivirus genomes with unique genomic islands were identified through clustering all genomes used in the initial analysis at 97% identity using mmseqs [54]. Genomes clustering together were then aligned using synteny plots from lovis4u [53] and manually inspected for unique genomic islands (Figure S9).

### Analysis of the potential origins of genomic islands in NCVs

To locate the potential origin of genes within the genomic islands of NCVs, all island proteins were subjected to a similarity search using BLASTP (e-value cutoff 1e-10) against the RefSeq database [66]. For each protein, the top non-viral hit was identified. The number of top hits to bacteria per island was normalized by the total number of genes on the island. The proportion of bacterial hits within the non-island region of the genomes was estimated in a similar manner (Figure 6a).

**Figure 6.**
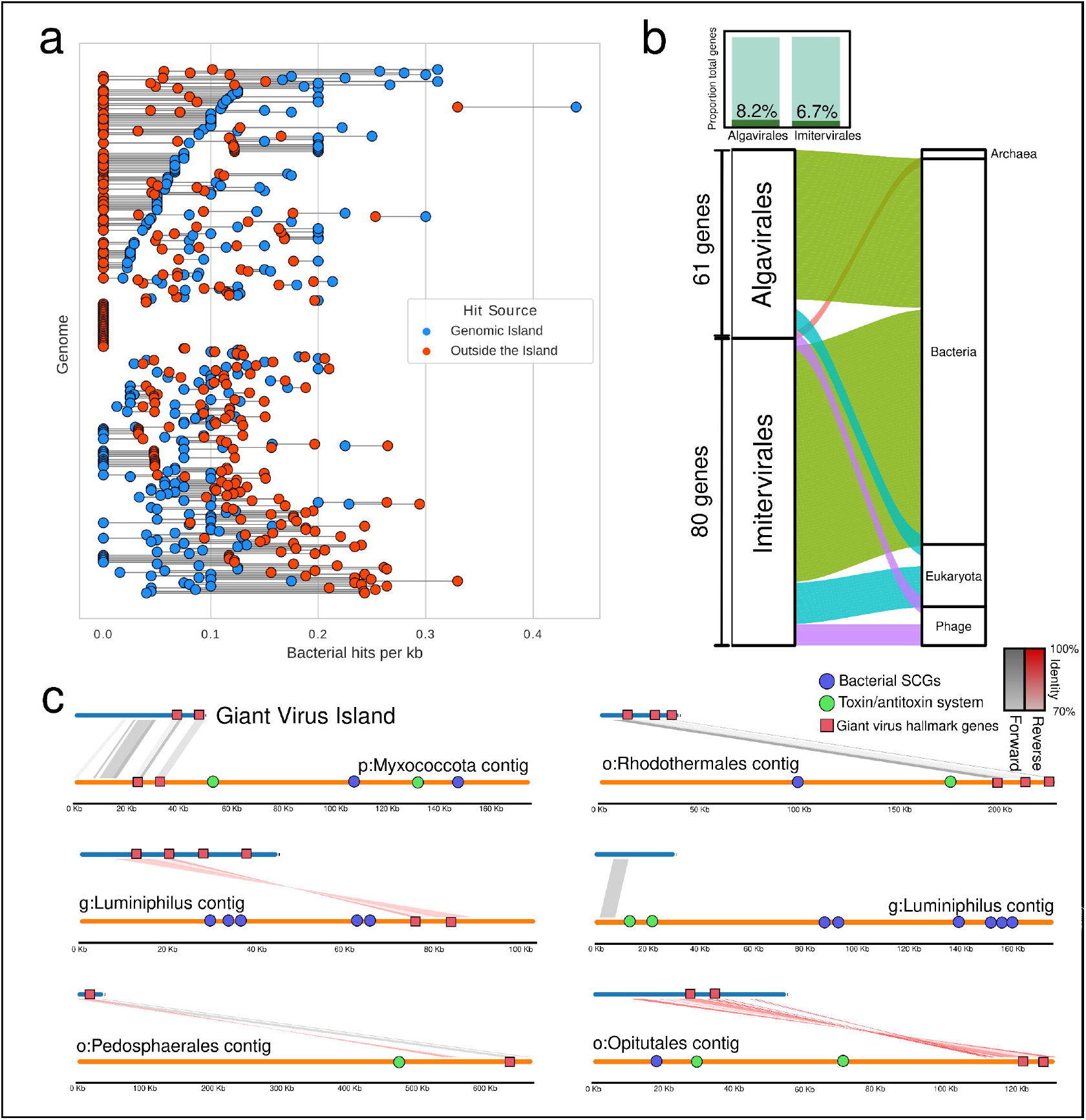
Evidence for the bacterial nature of certain giant virus genomic islands. (a) A plot showing the number of bacterial best hits (BLASTn) per kb inside of island regions (blue) and in the rest of the genome (red) of our identified giant viruses. The horizontal lines link points from the same genome. (b) Proteins from genomic islands from the San Francisco Estuary giant virus genomes were clustered with bacterial, eukaryotic, archaeal, and phage contigs from the same metagenome using mmseqs. The resulting alluvial plot shows the number of proteins from each giant virus order that clustered with protein from a cellular or phage contig (minimum 80% identity and 20% coverage). The proportion of total proteins that clustered with other (cellular) contigs in each order is shown in the barplot on top. (c) Total island regions from the San Francisco Estuary were also clustered with bacterial contigs from the same metagenome using mmseqs, resulting in 6 matches. The above genome maps highlight regions of synteny as well as giant virus hallmark genes (MCP, VLTF3), bacterial single-copy genes (SCGs), and genes involved in the toxin/antitoxin system. Bacterial contigs were classified based on consensus of their SCGs in anvi’o, and the lowest level of classification is given here. Zoomed-in synteny plots for selected islands with pfam annotations can be found in Figure S11.

To identify possible transfer of genomic regions across genomes of co-occurring NCVs and cellular organisms in the same environment, we leverage the metagenomic data from the San Francisco Estuary. First, all nanopore assembled contigs were filtered to a minimum length of 10 kbp. Then these contigs were classified to the kingdom level using a consensus approach of Tiara [67] and mmseqs taxonomy using the UniRef50 database [54], [68]. The proteins in the NCV genomic islands were clustered with a database of proteins from other contigs in the dataset using mmseqs linclust with a minimum sequence identity of 80% and a minimum coverage of 20%. Only island proteins that clustered with a protein on a contig with known taxonomy were further illustrated in the alluvial plot using ggplot2 (Figure 6b) [69].

To obtain further evidence of island transfer between NCVs and bacteria we identified entire island regions found in bacterial contigs. First, all bacterial contigs > 100 kbp were obtained from the previous analysis. The NCV islands were then aligned to these regions using BLASTn with a minimum alignment length of 25% of the island and an e-value cutoff of 1e-20 [61]. The resulting bacterial contigs were assigned taxonomy using single-copy genes in Anvi’o (anvi-estimate-taxonomy) [41]. Contigs that were found to have these islands were then binned with other contigs in the San Francisco long-read metagenome using Semibin2 [70], and symbiotic status of the bins was identified using symclatron [71] after manual bin refinement in Anvi’o [41]. Genome maps were made using Proksee [72].

To ensure that the island regions on the bacterial contigs were not erroneous due to chimeric assemblies, the nanopore long reads from the San Francisco Estuary were mapped back to the bacterial contigs using minimap2 to identify reads that spanned both the island regions and the flanking genomic regions [44]. The alignment results were then visualized using pysam and a custom script.

## Results and Discussion

### Genomic islands are widespread across NCV diversity

Utilizing our pipeline and 369 high-quality NCV genomes, we identified a total of 307 genomic islands with flanking direct repeats across 187 unique genomes (51%). Islands ranged from 5-135 kbp in length, and 68 genomes had more than one genomic island. Genomic island regions occupied an average of 15% of the genome of the virus that had them, with the highest percentage being 44% (Figure 1).

All 5 phylogenetic orders searched contained genomes with genomic islands and there did not appear to be phylogenetic enrichment of genomic islands within a particular order or family of NCVs (Figure S2). However, there tended to be more genomic islands per genome as genome size increased (R^2= 0.44) (Figure S3). Genomic islands also tended to have slightly higher GC % than the rest of the genome (by around 2%) with no noticeable trend in differences in codon bias or gene length (Figure S4). These findings align with what has been found in bacteria, as bacterial genomic islands tend to have GC differences ranging from 2-10% [73], [74].

Using the San Francisco Estuary short-read data for read recruitment, 53 of the 72 islands (74%) identified in the San Francisco Estuary NCV genomes were also metagenomic islands, showing deviations in read recruitment (+/- 2 SD) of island regions (Figure S1b). This finding supports the possibility that these islands can either be hypervariable regions within a viral population or widely distributed across many distinct viral populations, as has been shown elsewhere [32], [33], [75].

### Functional diversity and phylogenetic differences in genomic island proteins

A total of 12,691 proteins were found to be encoded within the genomic island regions, representing 6,894 unique protein clusters when clustered into orthologous groups. Of these, 42 proteins were significantly enriched in the islands of a particular NCV order (using a Fisher’s exact test) (Figure 2a). Many of the identified enriched proteins contained domains of unknown function according to PFAM, but further embedding-based homology searches (see methods) showed they were related to viral membranes and structure, highlighting unique mechanisms and proteins that may be used in capsid assembly and attachment between NCV orders [76].

Looking at key functional differences between NCV orders, Imitervirales genomic islands tend to encode more genes on average related to DNA repair and environmental response, such as MutS, rhodopsins, superoxide dismutase, and photolyase, while photosynthesis genes were found nearly exclusively in Algavirales islands, consistent with previous genome-wide findings [3], [77] (Figure 2b). Across all orders, islands encoded high proportions of genes with likely roles in host interaction, with the most common types being surface adhesion proteins, methyltransferases, and glycosyltransferases (Figure 2b) [78], [79]. Interestingly, multiple studies have shown that bacterial genomic islands frequently harbor genes involved in similar functions [80], [81], [82], and recent studies on hypervariable regions within the genomes of some prasinoviruses reported these functions as well [32]. These suggest some functional parallel between bacterial and NCV genomic islands, which might point to similar adaptive functions that are mediated through the genomic islands.

Clustering the genomic islands based on the presence/absence of unique orthogroups (see methods) revealed 9 clusters. (Figure S5). Of these 9 clusters, 7 formed tight groups, likely due to the close phylogenetic similarity of the genomes from which the islands originated. The largest cluster was characterized by the sporadic presence of numerous orthogroups, highlighting the high diversity of proteins found in most genomic islands. Further evidence for this high diversity comes from the fact that of the 12,691 island proteins, 5,390 are singletons (42.5%). This is consistent with previous observations in diverse bacterial genomic islands that harbor large amounts of hypothetical proteins [83], [84].

### Genomic islands in NCVs are enriched in surface adhesion proteins

The island proteins were mainly categorized as metabolic, interaction, or replication proteins (Figure 3). Of the 369 total islands, 75% had at least one protein involved in host interaction, 71% had a replication protein, 58% encoded a gene with possible metabolic functions, and 16% had all three types of genes encoded on the island (Figure 3ab).

To determine the functional specialization of the NCV genomic islands, we performed a functional enrichment analysis comparing genes within genomic islands vs those encoded in the rest of the genome. This revealed that genomic island regions were significantly enriched in various surface adhesion-related proteins (Ig-like domain, YadA, Collagen helix repeat, F5/8, Chaperone of endosialidase), as well as a gene related to DNA mobility with an HMG-box domain (Figure 3c).

Further phylogenetic investigation into the enriched surface adhesion proteins demonstrated that they seemed to be of bacterial origin, clustering within clades of similar proteins found in bacterial isolate genomes (Figure S6). Many of these enriched surface adhesion proteins have known adhesion roles within bacteria and phage communities, such as the YadA domain having a role in adhesion of bacteriophage tail fibers [85], as well as the cell-to-cell adhesion role of Ig domains, being the most widely used class of adhesion protein among viruses [86], [87](Figure 3d). Other proteins (F5/8 and FG-GAP) have been previously identified in genomic island regions of phages and NCVs infecting marine alga *Ostreococcus* [32], [88], highlighting the potential unique role these islands play in host adaptation through the exchange and capture of surface adhesion proteins.

While surface adhesion has not been an area of in-depth study within NCVs, it is undoubtedly an important step in successful infection, as most NCVs enter the host cell through phagocytosis or receptor-mediated endocytosis, both of which rely on some form of surface adhesion [10], [89]. One example of a well-studied surface adhesion protein in a NCV is the Mimivirus fibrils, which bind mannose and N-acetylglucosamine to mimic bacteria when binding to the surface of the host [90], [91]. Iridoviruses also encode IG-like domains, similar to those found to be enriched in our study, that have a predicted role in cell adhesion and evasion of the host immune system [92]. Within the realm of eukaryotic viruses, surface adhesion proteins are a key area of host adaptation, and studies have shown that switching of surface proteins is often indicative of host switching [93], [94]. Specific combinations of surface adhesion proteins have also been shown to be important for host entry, as is shown in HIV having both an IG-like domain, along with co-receptors to bypass multiple different cellular “gates” [95].

Our findings of enrichment of surface adhesion proteins within genomic island regions of NCVs are intriguing, as they show that these “keys” can be potentially shared throughout virus populations to aid the binding and opening of diverse cellular “locks”. This acquisition could allow NCVs to broaden their host range or counter changes in surface recognition proteins from their current host as part of the host-virus arms race, increasing viral fitness.

### Genomic islands can be widely shared throughout NCV population

Leveraging the long-read sequenced giant viral populations of San Francisco Estuary, we evaluated the population-level dynamics in genomic islands across closely related NCVs. Specifically, we identified a conserved island shared across 56 NCV genomes (order Algavirales) and contigs (Figure 4a). Mapping short reads from the same dataset to both the shared island region and the rest of the genomes revealed that the islands were, on average, ∼4x more abundant than the genomes they were derived from, providing further evidence for the exchange of these islands across related NCV populations (Figure 4c).

Further phylogenetic and ANI analyses showed that the 56 contigs and genomes sharing this island region formed at least two closely related but distinct clades within the order *Algavirales*, potentially representing genus-level divergence (Figure 4d, e; Figure S7). Across these genomes, the islands shared a broadly conserved gene repertoire, with many orthogroups present in both clades (Figure S8). However, the organization of genes (synteny) within the island did not strictly mirror viral phylogeny - in several cases, closely related genomes harbored islands with markedly different gene order, while showing greater structural similarity to islands found in genomes from the alternate clade (Figure 4b). This discordance between viral phylogeny and island architecture suggests that the evolutionary history of these regions is partly decoupled from that of the surrounding genome. One parsimonious explanation is horizontal transfer of the island region between related viruses, potentially during co-infection of the same host cell, although recombination and structural reshuffling within the island could also generate the observed mosaic patterns.

The shared islands contained many virion module proteins (minor and major capsid) as well as proteins putatively involved in viral replication and evasion of host defenses, such as methyltransferases and restriction enzymes (Figure 4d) [96], [97]. The widespread distribution of a viral surface antigen protein in these islands also gives evidence to the hypothesis that genomic islands can be beneficial to closely-related populations through the sharing of cellular “keys” for entry and successful infection.

### Genomic islands are a driver of hypervariability among closely related NCVs

While we showed that some genomic islands can be shared widely across a NCV population, these islands aren’t always totally identical, as genes can be added, deleted, or shuffled around within the island region, creating a hypervariable region in the genome that differs between even closely-related NCVs [32], [88]. To investigate this phenomenon, we examined several groups of closely related NCVs from diverse environments. Using our genomic island detection framework, we identified conserved island loci that act as modular regions of genomic variability. In these loci, islands share similar boundaries and portions of core gene content, but differ substantially in gene composition, organization, and sequence identity.

A clear example of this pattern occurs within prasinoviruses. Building on recent observations of hypervariable genomic regions in these viruses [32], we expanded the analysis by identifying 13 additional closely related genomes from the San Francisco Estuary that contain islands located at the same genomic loci. Although these islands share conserved flanking regions and portions of core gene content, their internal organization is highly variable, particularly in genes encoding surface adhesion proteins (Figure 5a). Overall, this analysis demonstrated that these island/hypervariable regions persist over ocean basins (previous prasinoviruses were isolated from the North Sea, Mediterranean Sea, and Pacific Ocean near Hawaii) with only minor changes to potentially adapt to different host regimes [32]. The conservation of this island region provides strong evidence that this island and those similar to it are critical to viral fitness and host adaptation [24].

This phenomenon of hypervariability within similar genomic islands isn’t unique to prasinoviruses, as we identified a similar case in closely related Mimivirus strains from different aquatic basins (Figure S9). For four different Mimivirus strains, we identified genomic islands (∼55 kbp each) with a well-conserved gene order structure in the first half of the island. However, this conservation falls apart in a hypervariable region of the island made up mostly of surface adhesion proteins such as the collagen helix repeats, adhesin autotransporters previously identified to be enriched in island regions (Figure 3c), and the YadA head domain. No two islands shared perfect synteny in these regions, and even similar proteins show differences in sequence and length.

In addition to hypervariability within islands, we also identified variability in terms of presence or absence of genomic islands in closely related genomes (Figure 5bc). Comparing two closely related Chlorella viruses (CVR-1 and OR0704) isolated from different water basins (Germany and Oregon) revealed a genomic island present in OR0704 that was missing in CVR-1 (Figure 5b). This island region contained many surface adhesion glycoproteins, common in Chlorella viruses, as well as a chaperone of endosialidase, which also has a putative surface adhesion role [98], [99].

A similar occurrence was also observed within two genomic islands from similar NCVs within the same ecosystem (Lake Biwa, Japan). One of these NCVs (LBGVMAG_0006) contained a large 76 kb genomic island that represents a major site of divergence from the otherwise syntenic LBGVMAG_0143 (Figure 5c). This genomic island contained a surface adhesion protein as well as other genes involved in replication (polymerases) and host adaptation through glycosyltransferases [79], [100]. Interestingly, this genomic island also contained a MutS gene, a commonly found DNA mismatch repair gene found in NCVs [3]. Upon further inspection of the data from the initial study, both of these viruses were predicted to occupy the same niche-layer of the lake (epilimnion), but the island-containing LBGVMAG_0006 was estimated to be 11.7x more abundant than the similar LBGVMAG_0143 [35]. This finding may suggest that the island region is critical to efficient viral infection and reproduction in this ecosystem, and the DNA repair capacity against UV damage or modulation of host adhesion might play a role in the success of this virus strain over the strain without this island. [101]. This example highlights the possible ecological advantages genomic islands likely confer to diverse NCVs.

Overall, these four examples demonstrate that hypervariability in genomic islands containing surface adhesion proteins may be a widespread phenomenon within NCVs and may be a key aspect of their environmental fitness in adapting to changing hosts or host-defense mechanisms. This hypervariability is likely driven by a combination of gene gain and loss as well as gene transfer between similar NCVs [19].

### Some genomic islands have bacterial origins

Given that genomic islands are widespread in NCVs and potentially aid in the virus-host arms race, we sought to identify the potential origins of these islands. It is well known that NCVs can exchange genes with their eukaryotic hosts as well as other viruses [19]. Interestingly, many NCVs are known to harbor genes of bacterial origin, but the driver or mechanism of transfer from bacteria remain unclear given that bacteria are not hosts of NCVs. In our study, we noticed that island regions were sometimes enriched in bacterial, rather than eukaryotic genes (Figure 6). Using a best-hit approach, we identified 137 islands (37%) that were enriched in bacterial hits per unit length (kbp) when compared to the non-island part of the same genome (Figure 6a).

To further investigate the potential for gene transfer between NCV genomic islands and other cellular organisms, we performed a protein clustering between genomic islands of NCVs from the San Fransisco estuary and other cellular contigs identified within the same metagenome data (Figure 6b). This clustering analysis revealed 141 proteins clustering both within the NCV islands and cellular contigs, with the majority clustering with bacterial contigs (113) (Figure 6b). These proteins came from both Imitervirales and Algavirales genomes in similar proportions (6.7 and 8.2%) and represent putative recent transfer events given the high similarity (>80%) to the cellular hits [102], [103]. Annotations of the clustered proteins revealed that the most frequent transfer events included the FG-GAP domain surface adhesion protein found in both Algavirales and Imitervirales genomes (Figure S10). Other surface adhesion proteins were also among the most transferred, including 2 chaperones of endosialidase and a Chlorovirus glycoprotein (Figure S10). As described earlier, many of these surface adhesion proteins were also found to be phylogenetically related to proteins from bacterial isolates (Figure S6).

In addition to evidence of many individual genes within genomic islands being of bacterial origin,, we also found evidence of partial or near-entire genomic islands to originate from bacterial genomes (Figure 6c, Figure S11). Within the SF estuary genomic islands, we identified six NCV islands that were also present (> 25% length coverage) on well-established bacterial contigs with taxonomic assignments. All of these contigs with established taxonomy could also be binned into bacterial MAGs (Figure S12). These contigs often contained multiple bacterial single-copy core genes used for taxonomy, as well as bacterial toxin/antitoxin systems (Figure 6c). To ensure these bacterial contigs were not assembly chimeras, nanopore long reads were mapped to the contigs, and reads were identified in each example that spanned both the island region and the rest of the contig (Figure S13). Of interest, there was even one read that spanned both a NCV major capsid protein (MCP) within the island and multiple bacterial genes, such as a gene with LTXXQ motif, ADP ribosylglycohydrolase, and S-adenosyl-L-homocysteine hydrolase (Figure S13). Bacterial contigs were classified into many different phyla, demonstrating that this phenomenon isn’t limited to specific phyla or orders.

While NCVs are thought to infect predominantly eukaryotic organisms [4], [7], gene exchange with other cellular organisms, such as bacteria, has been previously reported. In an overview of Chlorovirus gene transfer, Filee (2009) [104] found that 4-7% of Chlorovirus genes were bacterial in origin. Similar studies found chunks of genes within Mimivirus that seem to be transferred from bacteria near the termini of the genome or located near transposable elements [20], [21], [105]. There are also reports of three adjacent Mimivirus genes being potentially acquired from a *Clostridium* bacterium [20]. Our findings importantly suggest that not only do genomic islands in NCVs routinely harbor bacterial genes, but some islands feature large segments of DNA transferred from bacteria. Together, these results provide further evidence that gene exchange between NCVs and bacteria occurs and suggest that genomic islands may facilitate the transfer of large segments of bacterial DNA into NCV genomes, contributing to their genomic mosaicism.

### The Mechanism of transfer between NCVs and bacteria

The logical question remains as to how bacterial genes are transferred to NCVs that infect Eukaryotic organisms. Filee et al. (2007) [21] observed that two conditions would need to be met for gene exchange to occur (i) an ‘ecological niche bringing viral and bacterial DNA into close contact, and (ii) a mechanism to drive gene acquisition. We will explore both of these conditions in light of our findings.

The first condition is quite easy to satisfy, as the hosts of many NCVs are protists - complex, often single-celled, eukaryotic organisms that regularly host bacterial symbionts or phagocytose bacteria as a food source [106], [107]. These processes could create conditions where infecting viruses are introduced to foreign bacterial DNA inside the protist host, making it plausible for their integration into the genome. Of the six identified bacterial contigs containing NCV islands in our dataset, one of these was binned into a genome that was predicted to be an endosymbiote using symclatron (*Myxococcota* sp.) [71]. While the others were not predicted symbiotes, they include genera such as *Luminiphilus* that are known to be preferentially grazed by protists due to their large size and lack of defenses [108].

The second condition involves identifying a mechanism by which NCVs could acquire genetic material from a bacterium (or vice versa). It was noted by Filee et al (2007) [21] that recombination-primed replication could be an attractive mechanism for gene transfer as poxviruses and some Algavirales members undergo high levels of homologous recombination during replication [109], [110].

Another potential mechanism for the exchange of genomic islands between NCVs would be through mobile genetic elements (MGEs), genetic material that can move around a genome [111]. NCVs have previously been shown to contain a diverse array of MGEs with many having a putative prokaryotic origin [112]. We found that diverse MGEs are also prevalent within NCV genomic islands across all identified orders (Figure S14a), as 117 (38%) islands have at least one MGE (Figure S14b). Among the most common are restriction modification systems (R-M), restriction enzymes, and HNH endonucleases (Figure S14b). These R-M systems are of particular interest as they have been proposed to be another potential candidate for promoting recombination in NCVs, as is seen in herpesvirus recombination [113]. HNH endonucleases have also been previously identified in Algaviruses and Mimiviruses, often associated with other genes of bacterial origin [21], [114]. Some islands also appear to contain transposons within them, characterized by diverse transposases found within the island region (Figure S15).

Overall, these results suggest that certain NCV genomic islands were found to be exchanged with bacterial contigs and contain many MGEs to facilitate this transfer, supporting hypotheses generated by Filee el al. (2007) [21].

## Conclusion

Our study establishes genomic islands as a widespread and dynamic feature of giant virus genomes rather than rare anomalies of horizontal gene transfer. Across 369 high-quality genomes we identified 307 genomic islands in 187 viruses, showing that these regions are broadly distributed across Nucleocytoviricota and can occupy substantial portions of the viral genome. Their presence across all major phylogenetic orders indicates that island acquisition and turnover are recurring processes that shape the genomic function and host-virus co-evolutionary arms race. Many islands are enriched in genes associated with surface adhesion and host interaction, and several display signatures of hypervariability among closely related viral strains. These patterns indicate that genomic islands may act as flexible genomic modules where gene gain, loss, and rearrangement occur rapidly. Thus, our study establishes genomic islands to be a crucial component of giant virus genomes which possibly allow giant viruses to explore new infection strategies or host associations without restructuring the broader genome architecture.

We also revealed extensive gene exchange associated with these islands. A substantial fraction are enriched in genes of bacterial origin, with several cases of large genomic regions from bacteria to be transferred as GIs. These patterns suggest that genomic islands may participate in a broader network of genetic exchange occurring within infected protist cells, where giant viruses, their eukaryotic hosts, and associated bacteria interact within the same cellular environment. Importantly, our data also suggests the possibility of entire islands to be exchanged between closely related giant viruses supported by our synteny analysis and observation of the presence-absence patterns of entire islands within closely related strains. Together, these results are consistent with the idea that protist hosts can act as ecological hubs where genetic material from diverse biological partners encounters opportunities for exchange.

Future work will be needed to determine the functional consequences of individual islands and the mechanisms that mobilize them. Experimental validation of candidate adhesion proteins, for example, could clarify whether islands serve as hotspots for viral innovation in host recognition and entry. More broadly, integrating genomic islands into studies of giant virus ecology and evolution provides a framework for investigating how these viruses acquire, reorganize, and deploy genetic material within complex microbial communities, while also offering insight into why certain viral strains become ecologically successful whereas others do not, and how the acquisition and diversification of genomic islands may influence competitive dynamics among co-occurring viruses.

Together, our results suggest that the mosaic genomes of giant viruses are shaped not only by gradual gene accumulation but also by the repeated incorporation and remodeling of large genomic segments. Genomic islands therefore represent a major source of genome plasticity in giant viruses and provide an entry point for understanding how gene flow across diverse biological partners contributes to the evolution of the giant viruses in the biosphere.

## Acknowledgments

This work was supported by the Rosenstiel School of Marine, Atmospheric, and Earth Sciences, University of Miami. We would also like to thank Dr. Torben Nielsen and Dr. Lauren Lui (Lawrence Berkeley National Laboratory, Berkeley, California, USA) for their provision of the San Francisco Estuary metagenomic dataset and for help in editing the final manuscript.

## Data Availability

The code for recovering genomic islands is publicly available on github (https://github.com/BenMinch/IslaMine). All genomes used in the analysis as well as raw files for phylogenetic trees, alignments, genomic islands, and protein annotations are provided on FigShare (https://figshare.com/projects/_b_Widespread_genomic_islands_in_giant_viruses_shape_genome_plasticity_and_mosaicism_b_/273924).

## Figure Legends

