## Supplementary File for "Widespread Genomic Islands in Giant Viruses Shape Genome Plasticity and Mosaicism"

Supplementary Material for “Widespread genomic islands in giant viruses shape genome plasticity and mosaicism”.

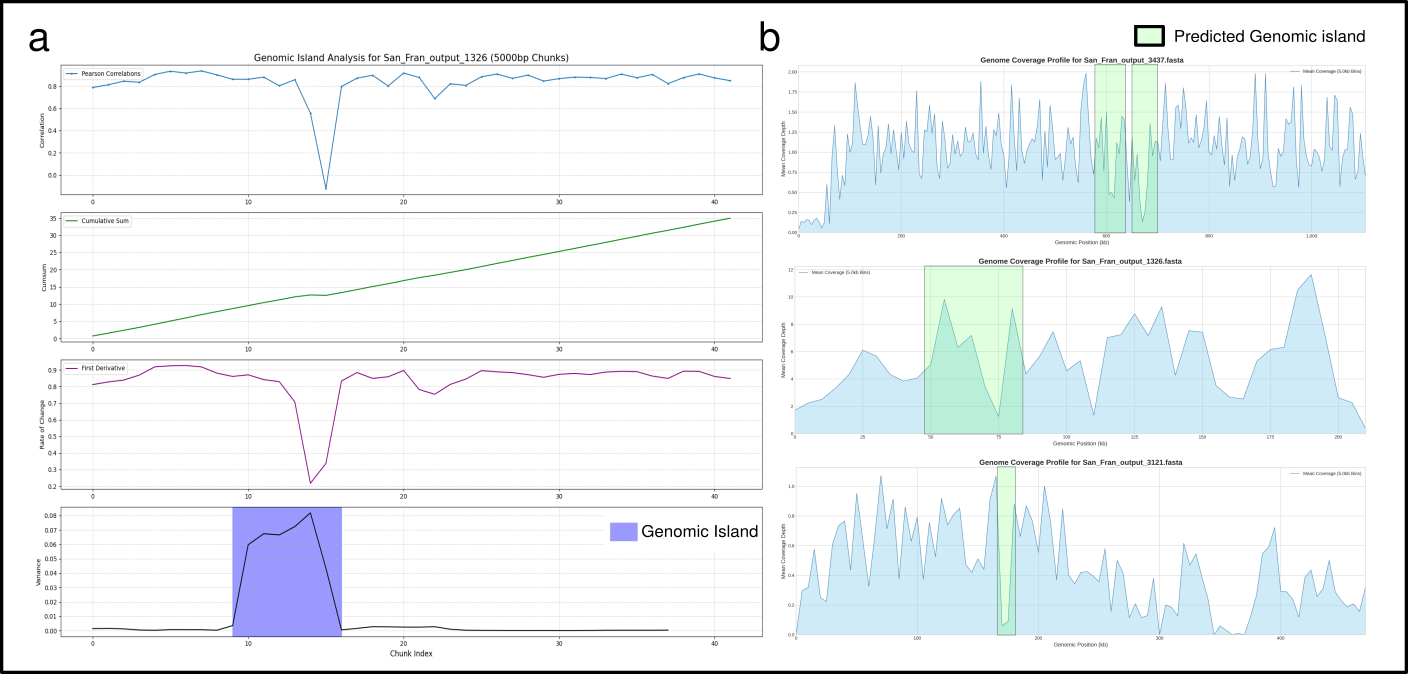

**Figure S1. Identification of genomic and metagenomic islands.** **(a)** A representative output of our genomic island identification algorithm. The panels show Pearson correlation of tetranucleotide frequency over 5 kb chunks of the genome, as well as the cumulative sum of these correlations and the first derivative. The final identified island is shown in blue. **(b)** Three representative giant virus genomes from the San Francisco estuary show that some of the predicted genomic islands are also metagenomic islands. Short reads were mapped to each genome separately at 95% identity, and coverage was visualized as mean depth in 5 kb bins.

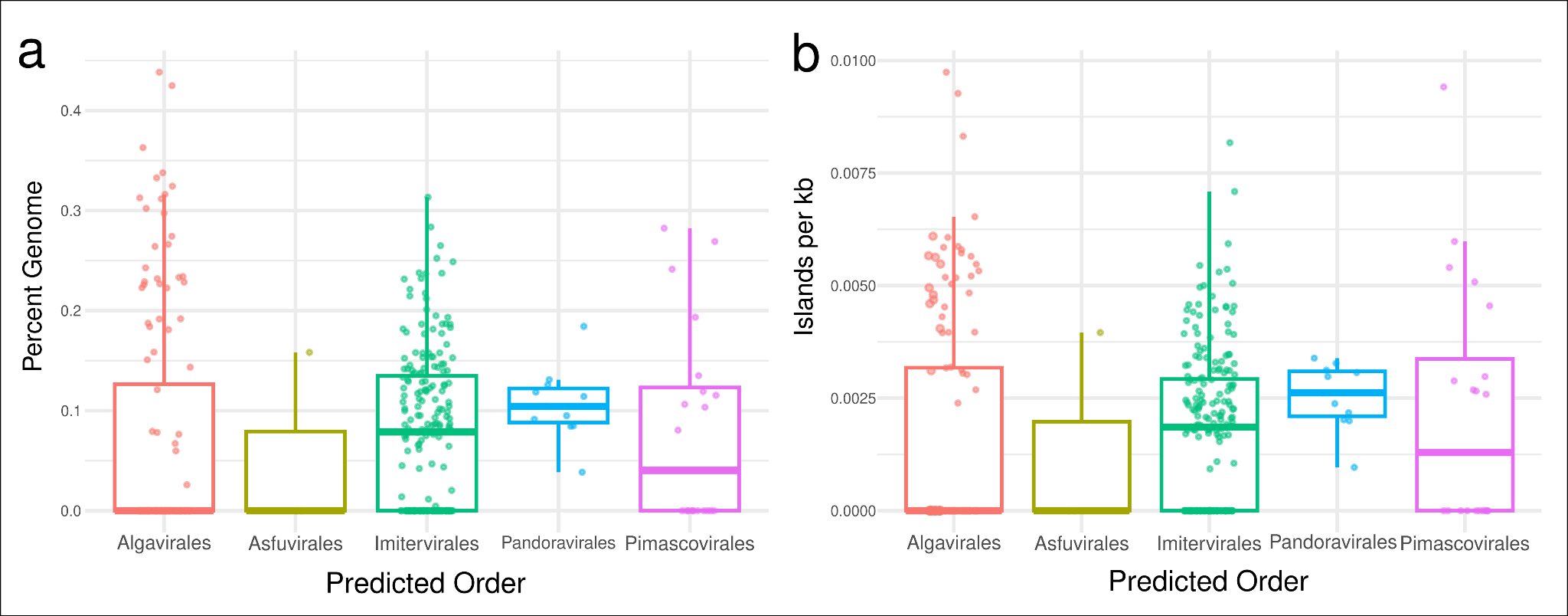

**Figure S2. Genomic island landscape across giant virus phylogenetic orders.** For each predicted giant virus order, the **(a)** percent of the viral genome composed of genomic islands and **(b)** the number of islands normalized to virus genome size (islands per kb) are shown as boxplots.

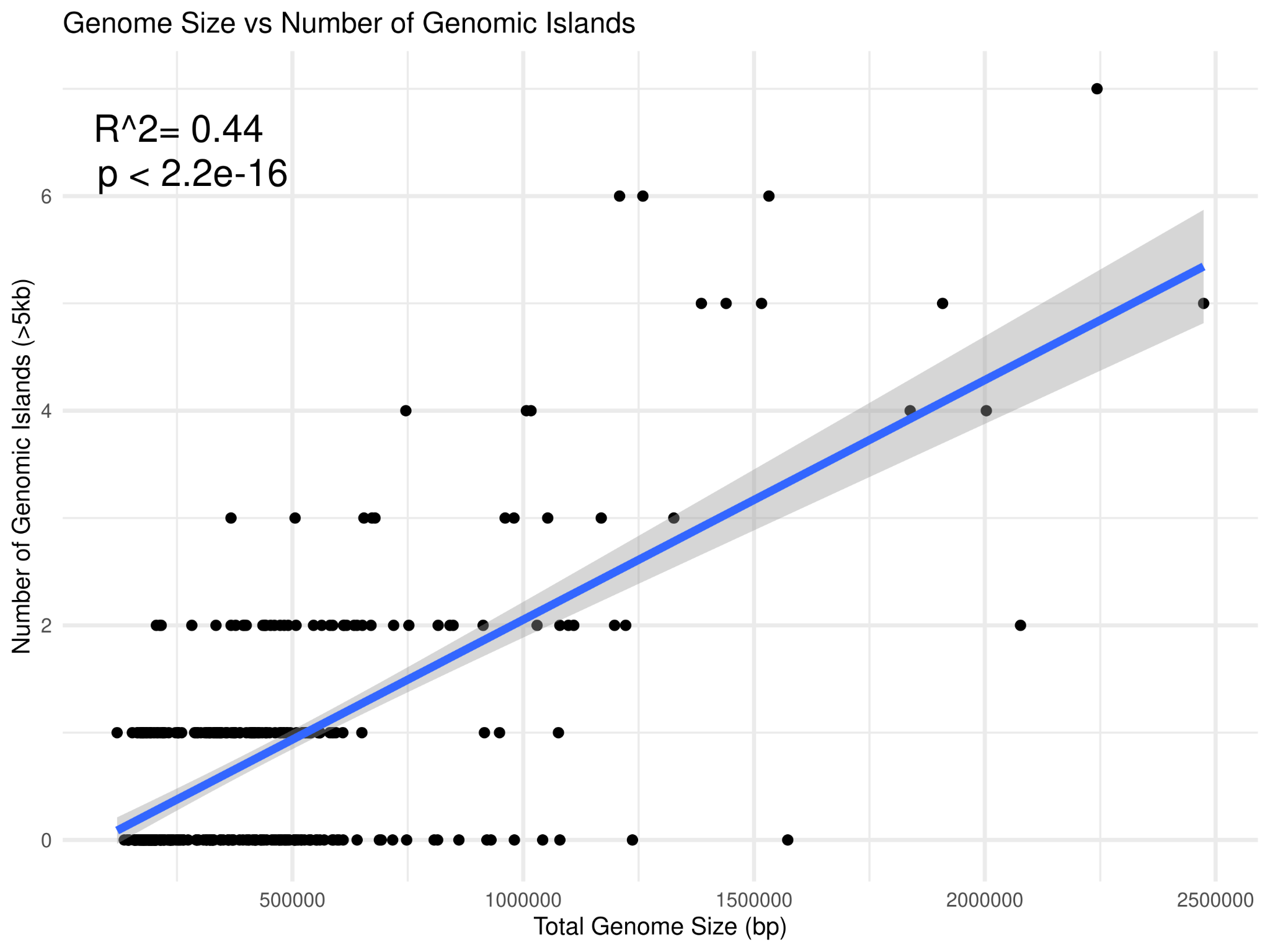

**Figure S3. Correlation between genome size and total number of identified genomic islands.** A scatterplot showing the number of genomic islands and total genome size. A linear regression was performed and is displayed on the plot.

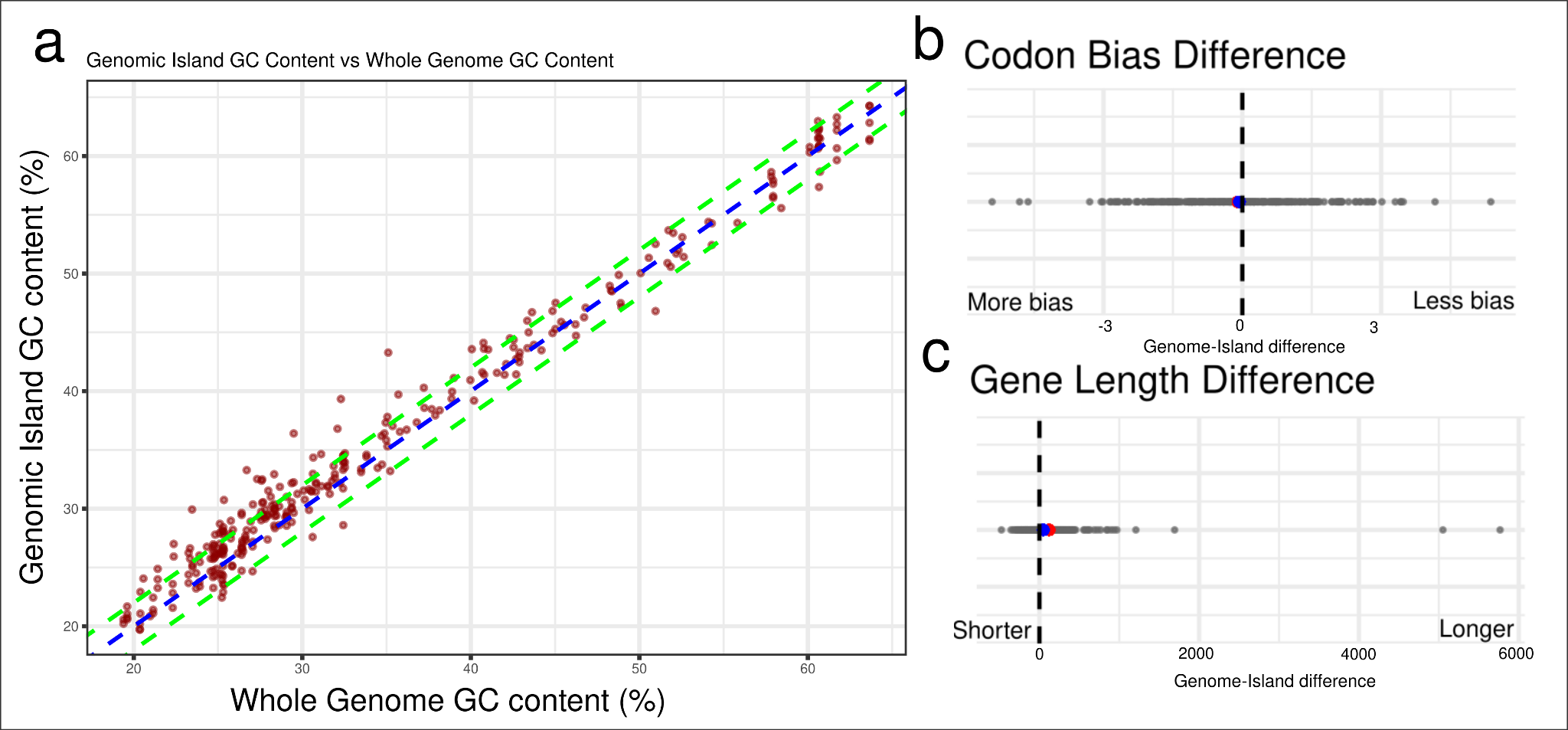

**Figure S4. Genomic characteristics of island regions in giant viruses.** **(a)** A comparison between the GC percentage of genomic islands vs the genome where they originated. The blue dotted line represents a 1:1 correlation while the green line represents a GC difference of +/- 2%. **(b)** Codon bias and **(c)** gene length differences between genomic islands and the rest of the genome. 1:1 correlation is represented by the dotted line in the middle, along with dots representing the average (blue) and median (red) values for genomic islands.

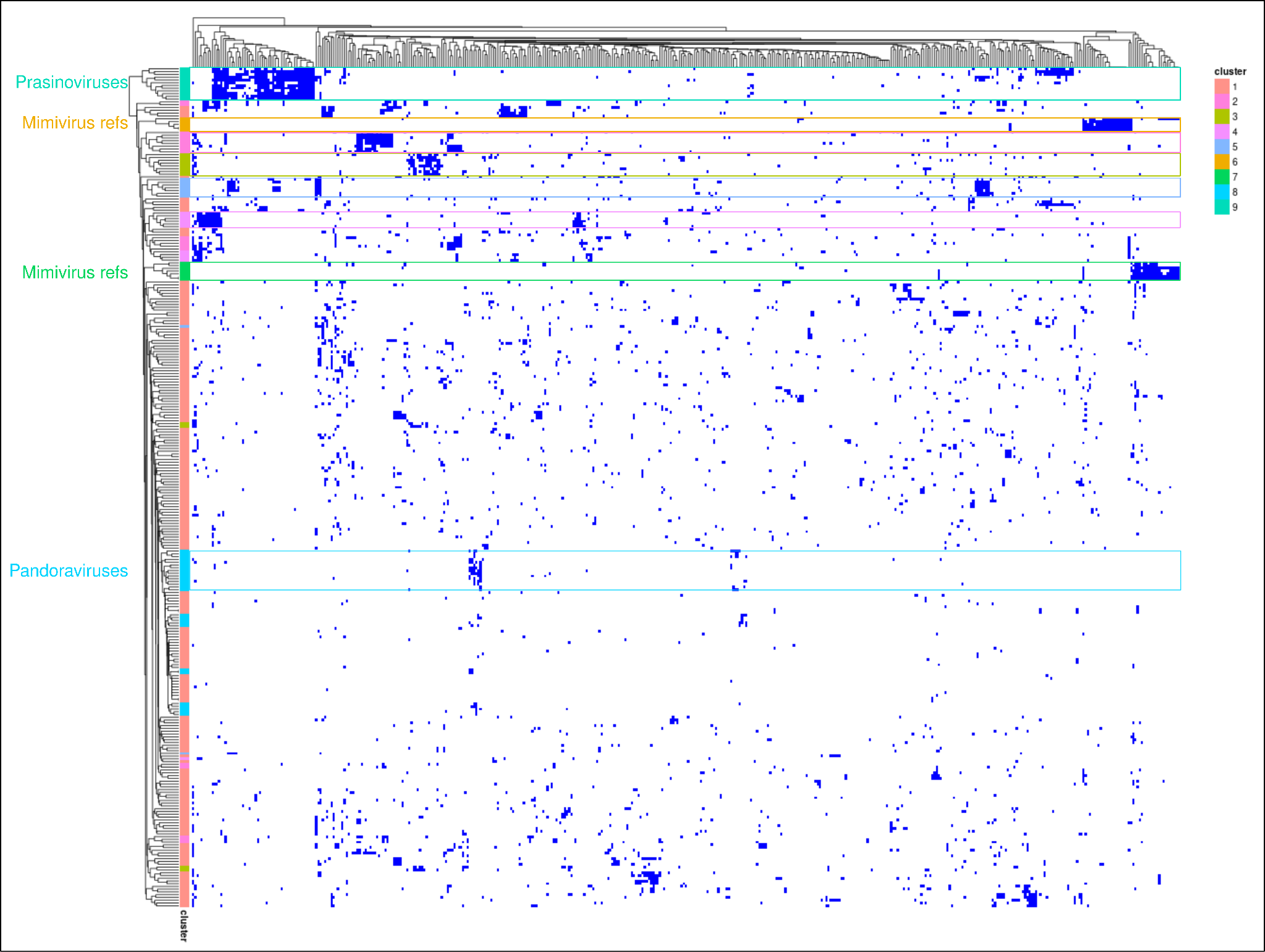

**Figure S5. Orthogroup clustering of genomic island proteins.** A heatmap showing the presence/absence of the 12,691 orthogroups identified through mmseqs clustering. Different clusters of islands were identified using hierarchical clustering with a minimum Davies-Bouldin index, and those clusters are shown in the color strip, with clusters showing close phylogenetic relationships being highlighted (Prasinoviruses, Pandoraviruses, and Mimiviruses).

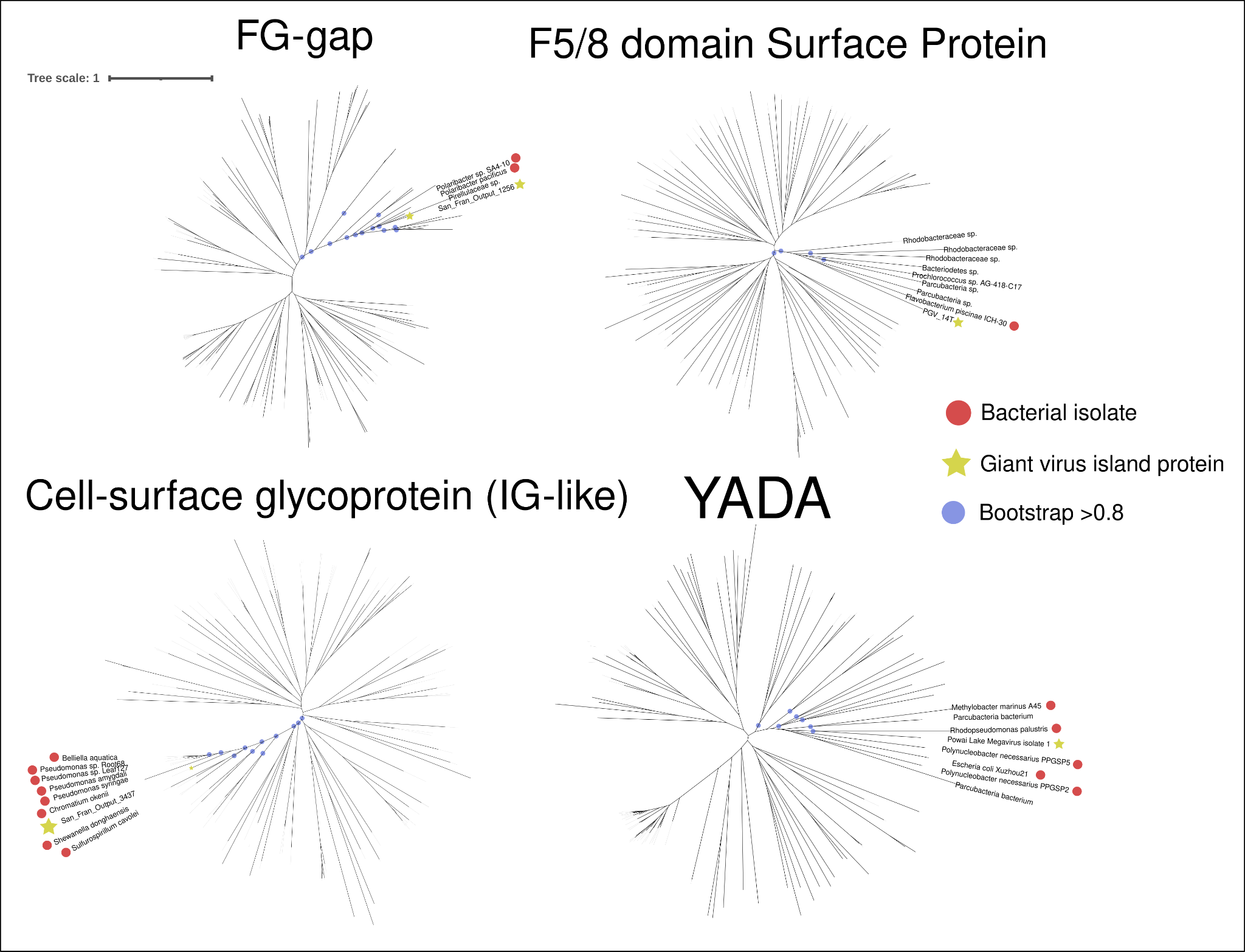

**Figure S6. Phylogenetic placement of genomic island-enriched surface adhesion proteins.** Phylogenetic trees for the most enriched surface adhesion proteins were made using a representative sequence from a giant virus genomic island and 100 top reference sequences from the Open Genomes Database within Seqhub. The trees were built using IQTREE and displayed using iTOL. Reference sequences representing bacterial isolates are highlighted with a red dot, and nodes with > 0.8 bootstrap support are also highlighted.

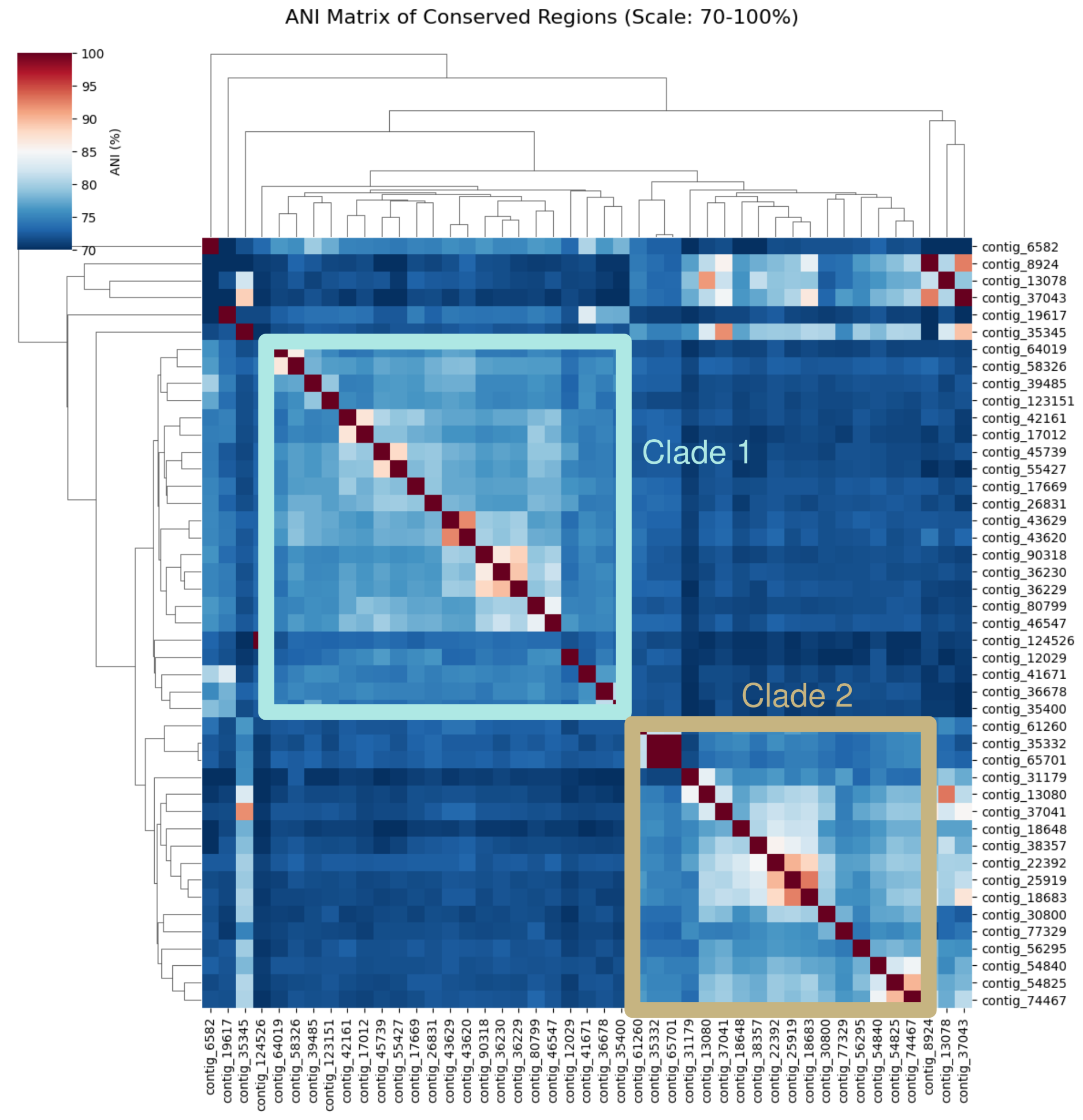

**Figure S7 ANI clustering of contigs with shared genomic islands.** All-vs-all average nucleotide identity (ANI) clustering was performed using PyANI, using the contigs containing shared island regions identified in Figure 4.

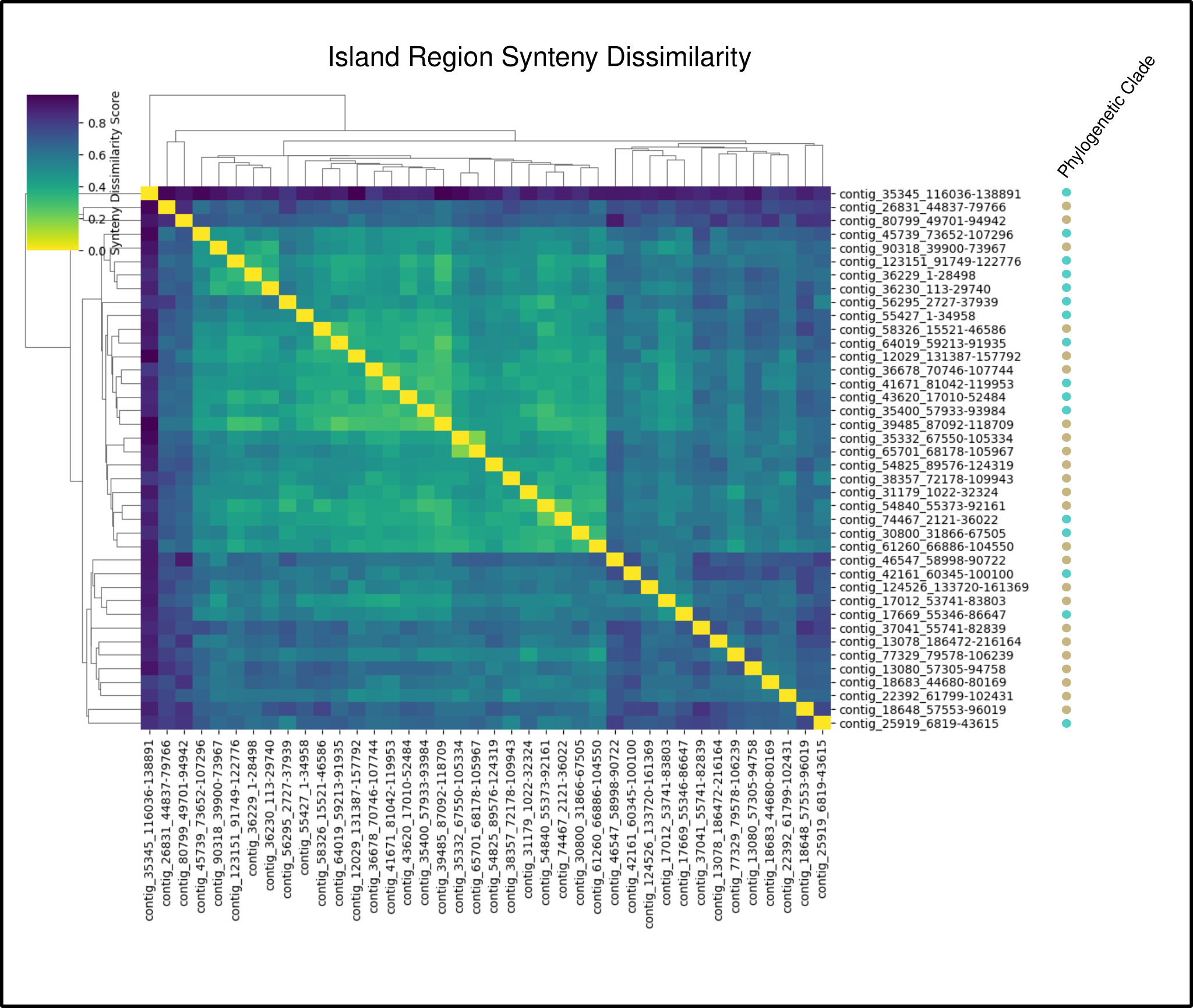

**Figure S8. San Francisco Estuary island synteny dissimilarity.** For each island identified in Figure 4, proteins were clustered into orthologous groups using proteinortho. The dissimilarity in the synteny between islands was calculated using the Levenshtein distance between islands, treating each orthogroup as a unique “word” in a sentence.

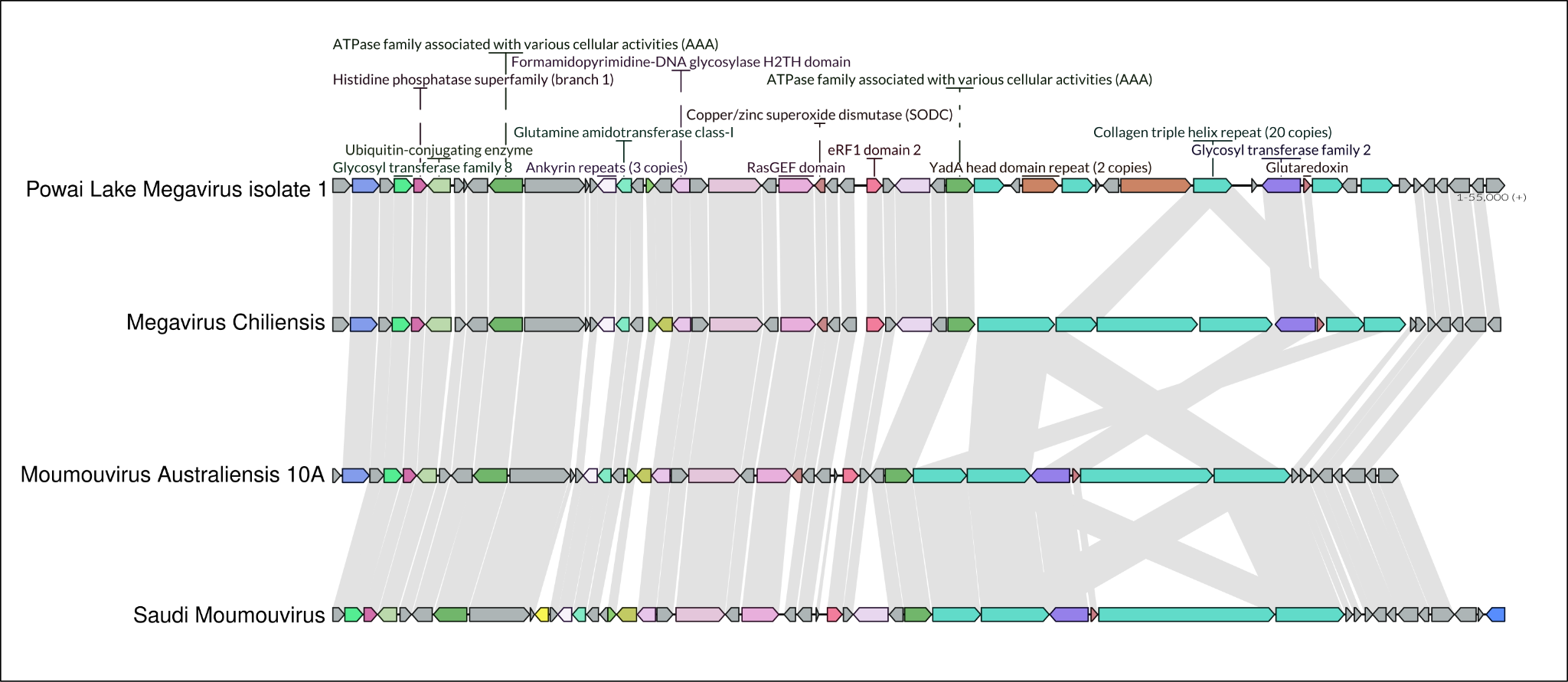

**Figure S9. Genomic island variability in similar Mimivirus strains from diverse aquatic basins**. A genome map of four genomic islands from similar Mimivirus genomes was constructed using lovis4u. Annotations were performed using the Pfam database, and synteny lines represent proteins with > 90% similarity.

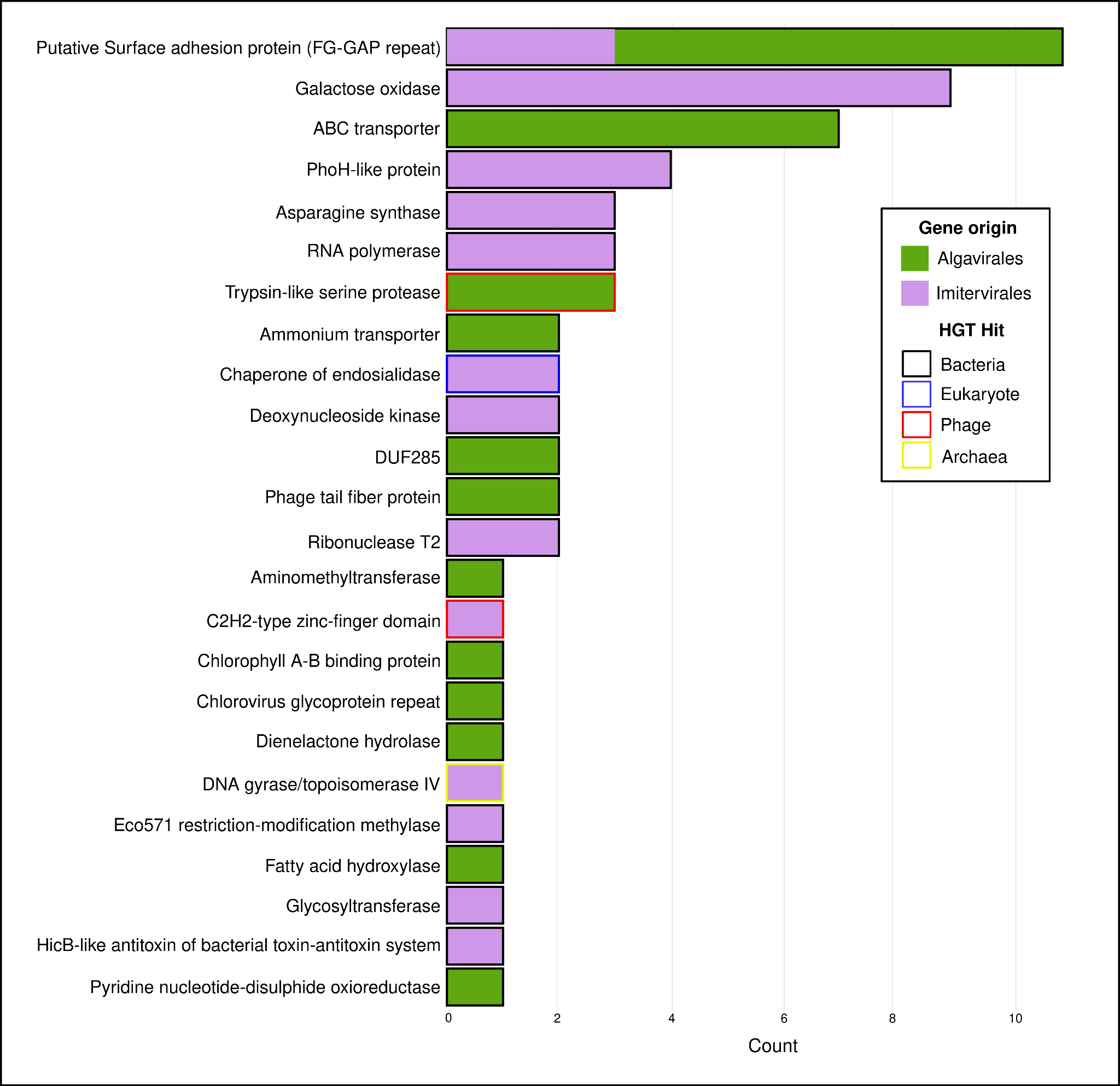

**Figure S10. The most commonly horizontally transferred genes between San Francisco Estuary giant viruses and other organisms**. Proteins from San Francisco giant virus genomic islands were clustered with cellular and phage proteins from contigs retrieved from the same metagenome (80% similarity threshold). The top 25 most transferred genes between these groups are shown here, along with the source of the protein (Imitervirales or Algavirales) and the contig where a similar protein was found.

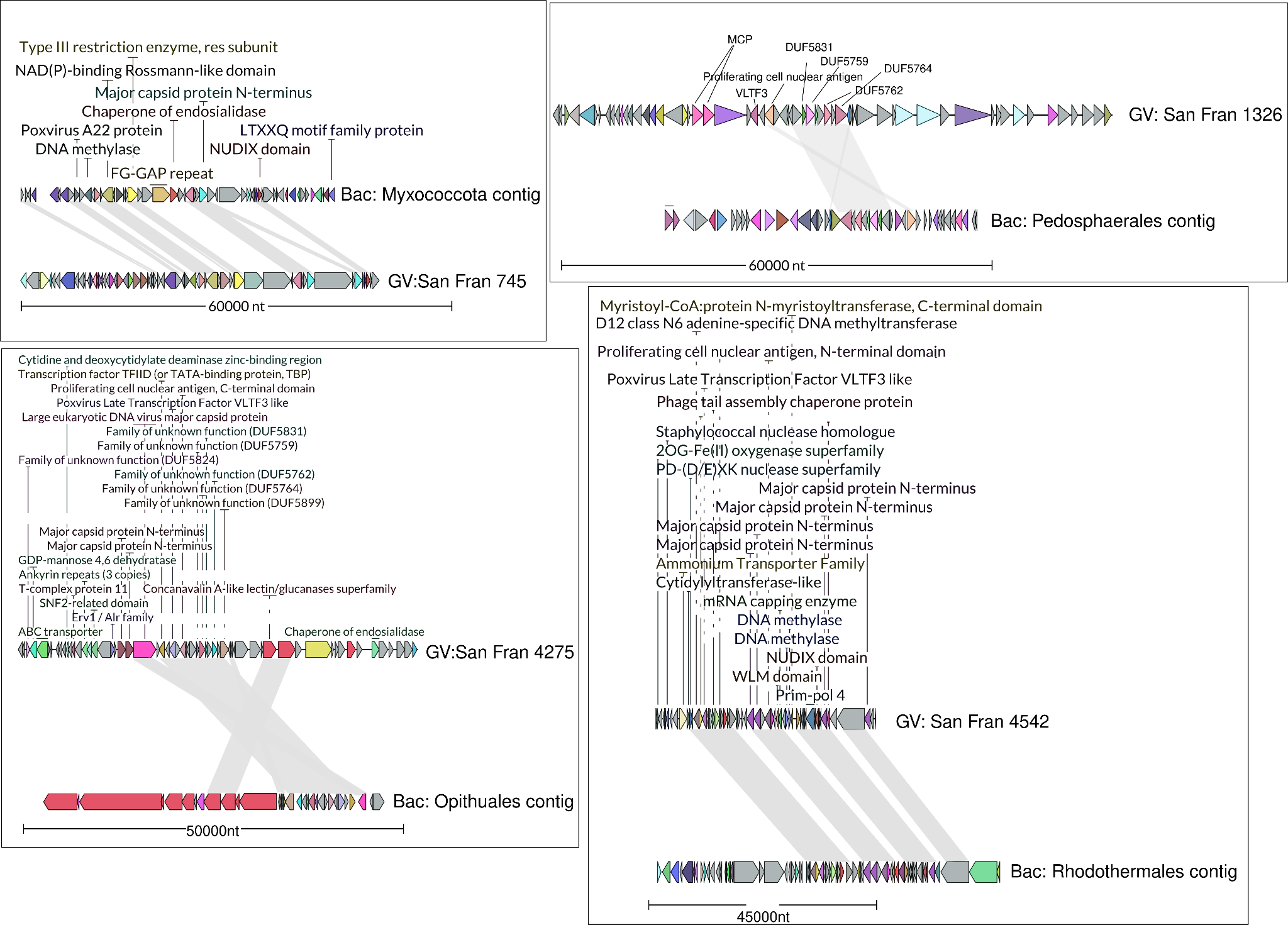

**Figure S11. Selected synteny plots for islands shared across giant virus and bacterial contigs.** In-depth synteny plots showing Pfam annotations for selected islands from Figure 6c. Plots were made using lovis4u.

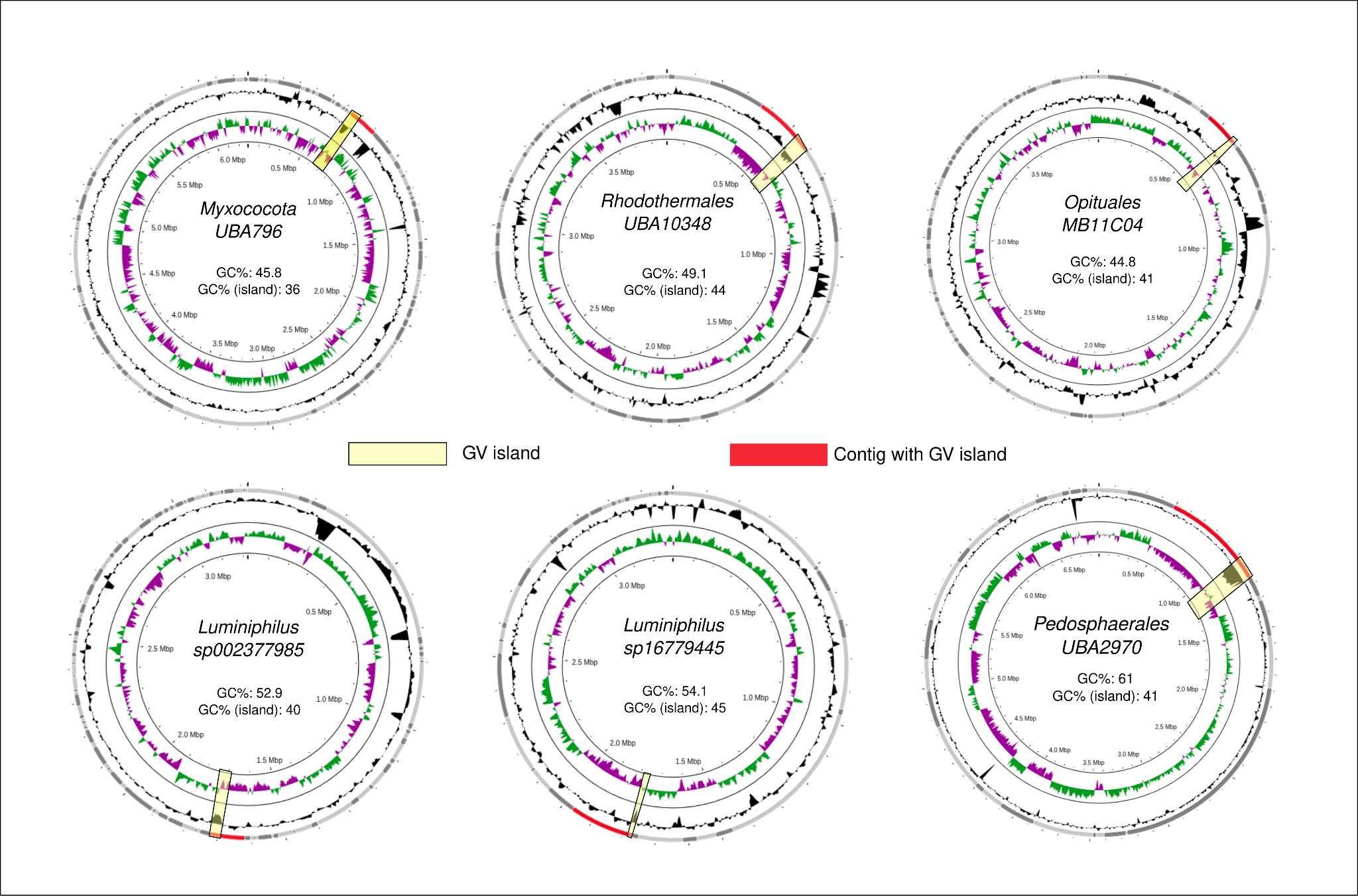

**Figure S12. Bacterial MAGs containing giant virus genomic islands.** Bacterial MAGs were recovered from the San Francisco Estuary long-read metagenome using Semibin2. Bins containing contigs of interest (Figure 6c) were further refined using Anvi’o to remove contamination. Genome plots were made using the Proksee web server, and contigs containing the giant virus island are highlighted. The inner ring is GC skew, and the outer is GC content, with the GC content difference between the genome and island region highlighted in the middle. Taxonomy was identified using anvi-estimate-taxonomy with bacterial single-copy genes (SCGs) present in the bins.

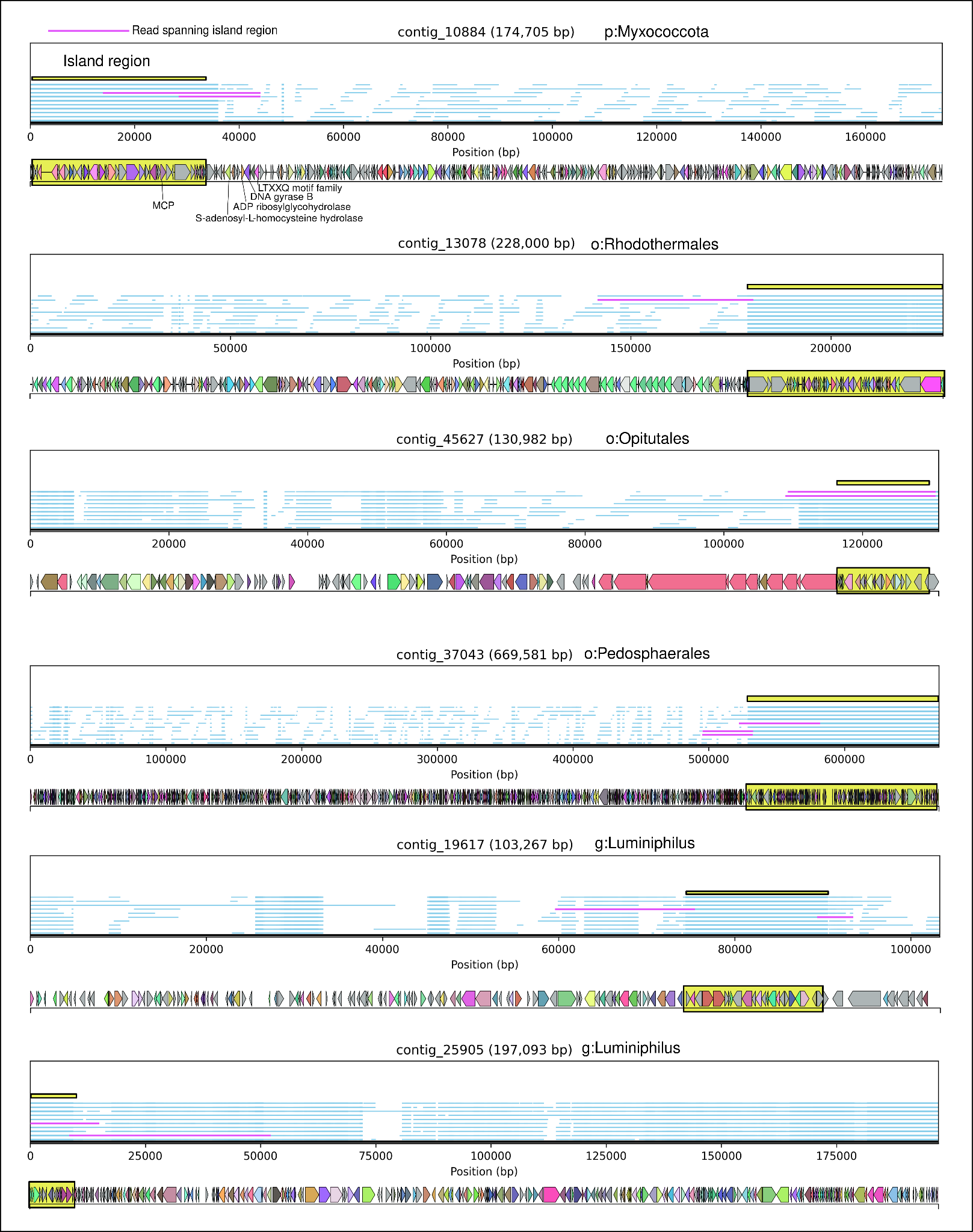

**Figure S13. Long read coverage of the bacterial contigs containing giant virus genomic islands.** To ensure against chimeric assembled contigs, long reads were mapped to bacterial contigs containing the giant virus genomic islands at 95% identity. The resulting mappings are visualized with each blue line representing individual reads. The island regions of these contigs are highlighted in yellow, and reads spanning both the island and the rest of the contig are highlighted in pink. Genome maps were made using lovis4u.

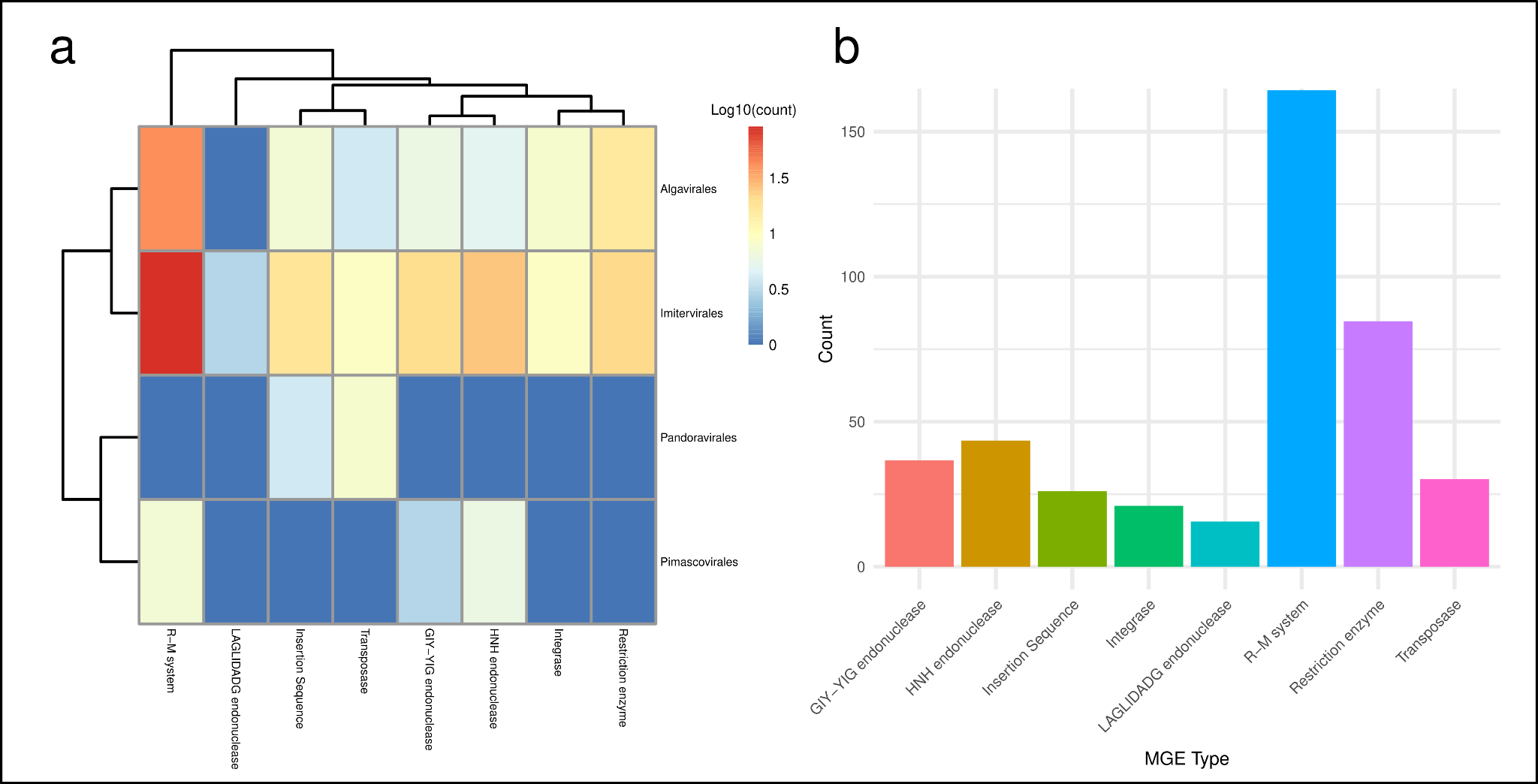

**Figure S14. Mobile genetic elements within giant virus genomic islands. (a)** A heatmap showing the abundance of different mobile genetic elements (MGEs) found within giant virus islands of different phylogenetic orders. **(b)** A barplot depicting the total counts of each MGE type found across all giant virus islands. MGEs were identified using REBASE, ISfinder, and the PFAM database.

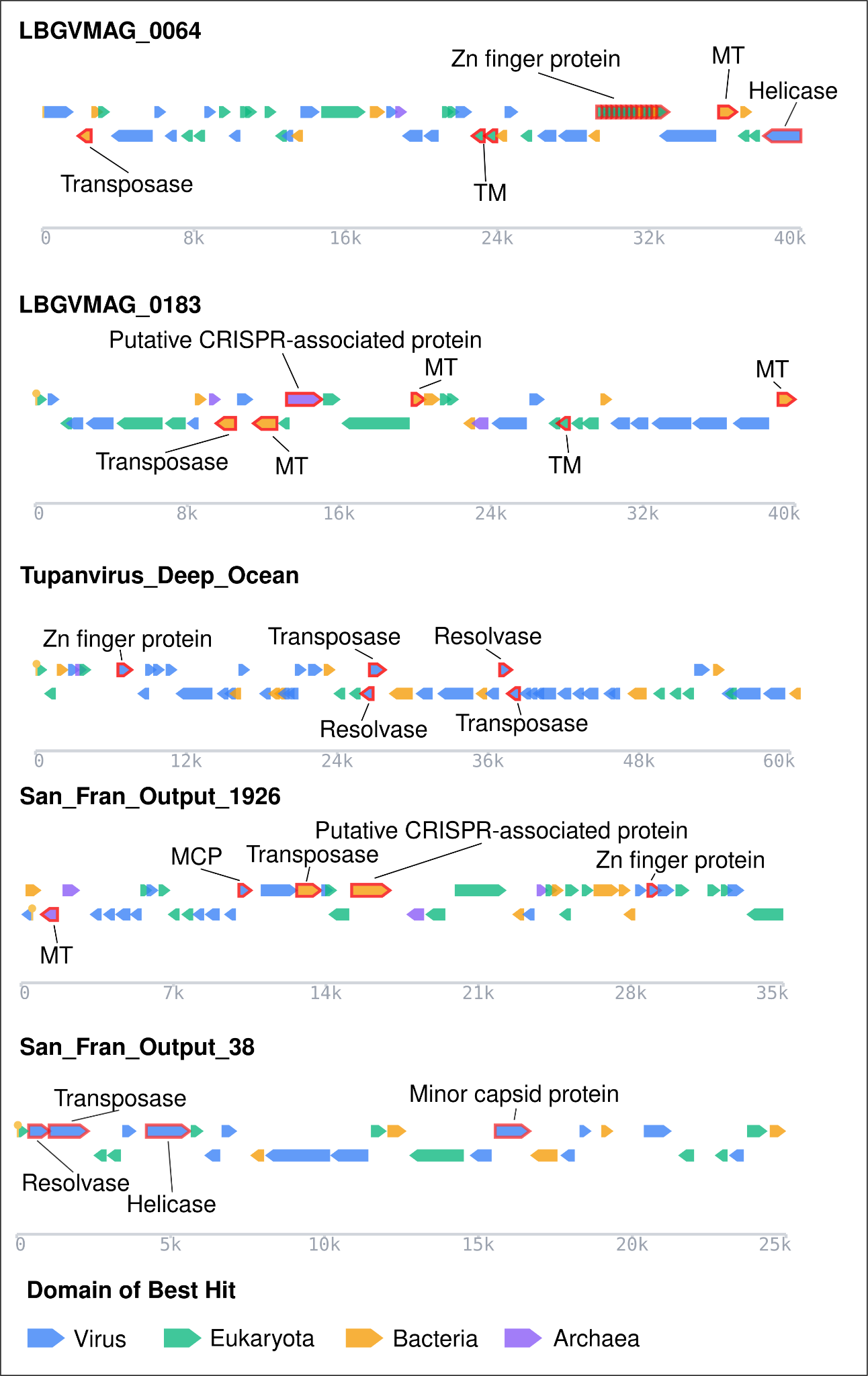

**Figure S15. Genomic islands containing transposases.** Several representative genomic islands containing transposases were mapped using Seqhub. Gene abbreviations are as follows: Methyltransferase (MT), Transmembrane protein (TM), Major capsid protein (MCP). The domain best hit was also obtained from the Seqhub platform based on the closest Swissprot hit.

| Virus | Reported Island Range | Recovered Island Range | Reference |
| --- | --- | --- | --- |
| GV1_CRY1 | 425-452 kb | 416-448 kb | Mukherjee et al., 2025 |
| EhV86 | 237-258 kb | 225-270 kb | Pagarete et al., 2014 |
| MCV-20T | 42-75 kb | 54-70 kb | Thomy et al., 2025 |
| OtV-06-1 | 42-75 kb | 44-64 kb | Thomy et al., 2025 |
| OtV-06-12 | 42-75 kb | 50-80 kb | Thomy et al., 2025 |
| OtV-06-4 | 42-75 kb | 30-75 kb | Thomy et al., 2025 |
| OtV-09-556 | 42-75 kb | 30-70 kb | Thomy et al., 2025 |
| OtV-09-557 | 42-75 kb | 30-70 kb | Thomy et al., 2025 |
| OtV-09-559 | 42-75 kb | 30-70 kb | Thomy et al., 2025 |
| OtV-09-565 | 42-75 kb | 60-72 kb | Thomy et al., 2025 |
| OtV-09-570 | 42-75 kb | 30-75 kb | Thomy et al., 2025 |
| OtV-09-573 | 42-75 kb | 30-70 kb | Thomy et al., 2025 |
| OtV-09-578 | 42-75 kb | 56-66 kb | Thomy et al., 2025 |
| OtV-09-582 | 42-75 kb | 30-70 kb | Thomy et al., 2025 |
| OtV-09-585 | 42-75 kb | 30-75 kb | Thomy et al., 2025 |
| OtV-19-O | 42-75 kb | 44-64 kb | Thomy et al., 2025 |
| OtV-19-P | 42-75 kb | 46-66 kb | Thomy et al., 2025 |
| OtV-19-R | 42-75 kb | 30-75 kb | Thomy et al., 2025 |
| OtV-19-T1 | 42-75 kb | 44-64 kb | Thomy et al., 2025 |
| OtV-19-T2 | 42-75 kb | 42-76 kb | Thomy et al., 2025 |
| OtV-Sylt2-5 | 42-75 kb | 30-70 kb | Thomy et al., 2025 |

**Supplemental Table 1. Benchmarking our genomic island recovery method against published giant virus genomes with identified genomic islands.**

| Category | Description | Example Genes |
| --- | --- | --- |
| Interaction | Genes putatively involved in cell-cell adhesion or viral interaction with host proteins. | Surface adhesion proteins, glycosyltransferases, and methyltransferases. |
| Environment Response | Genes putatively involved in a response to an environmental stimulus (UV radiation, light, heat, etc.). | Heat shock protein, Ice nucleation protein, MutS, rhodopsin, SOD. |
| Structure | Genes involved in viral structure and capsid formation. | MCP, mCP, A32 ATPase, VLTF3 |
| Replication | Genes involved in viral replication. | PolB, RNAPL/RNAPS, TFIIB, DNA topoisomerase |
| Metabolism | Genes involved in cellular metabolic pathways. | Citrate synthase, aconitase, cytodyltransferase |
| Unknown | Genes not categorized. | DUFs, small repeats |

**Table S2. Functional categories of genes in genomic islands.**
